# Deacetylation by sirtuins is important for *Aspergillus fumigatus* pathogenesis and virulence

**DOI:** 10.1101/2023.09.25.558961

**Authors:** Natália Sayuri Wassano, Gabriela Bassi da Silva, Artur Honorato Reis, Jaqueline A. Gerhardt, Everton P. Antoniel, Daniel Akiyama, Caroline Patini Rezende, Leandro Xavier Neves, Elton Vasconcelos, Fernanda L. Figueiredo, Fausto Almeida, Patrícia Alves de Castro, Camila Figueiredo Pinzan, Gustavo H. Goldman, Adriana F. P. Leme, Taicia P. Fill, Nilmar S. Moretti, André Damasio

## Abstract

Protein acetylation is a crucial post-translational modification that controls gene expression and a variety of biological processes. Sirtuins, a prominent class of NAD^+^-dependent lysine deacetylases, serve as key regulators of protein acetylation and gene expression in eukaryotes. In this study, six single knockout strains of fungal pathogen *Aspergillus fumigatus* were constructed, in addition to a strain lacking all predicted sirtuins (SIRTKO). Phenotypic assays suggest that sirtuins are involved in cell wall integrity, secondary metabolite production, thermotolerance, and virulence. *AfsirE* deletion resulted in attenuation of virulence, as demonstrated in murine and *Galleria* infection models. The absence of AfSirE leads to altered acetylation status of proteins, including histones and non-histones, resulting in significant changes in the expression of genes associated with secondary metabolism, cell wall biosynthesis, and virulence factors. These findings encourage testing sirtuin inhibitors as potential therapeutic strategies to combat *A. fumigatus* infections or in combination therapy with available antifungals.

## INTRODUCTION

Humans are daily exposed to the opportunistic fungal pathogen *Aspergillus fumigatus*^1^. Although the immune system of healthy individuals is effective in eliminating this microorganism, immunosuppressed patients are at high risk of developing invasive pulmonary aspergillosis (IPA), a disease with a high mortality rate^2^. Moreover, the emergence of COVID-19-associated IPA and antifungal-resistant isolates has raised serious medical concerns^3,4^. IPA pathogenesis depends on multiple host factors such as the immune system status^1,5–7^. Moreover, some characteristics of *A. fumigatus* are essential factors for full virulence, such as (i) conidial melanin and hydrophobicity that protect against reactive oxygen species (ROS) and damage by immune cells^8,9^, (ii) the ability to survive at high temperature^10^, and (iii) production of secondary metabolites (SMs), such as siderophores and gliotoxin^11,12^.

Previous studies indicate that chromatin conformation is involved in fungal virulence and host-pathogen interactions^13^. Moreover, post-translational modifications (PTMs) that occur in histones and non-histone proteins, such as acetylation, methylation, and phosphorylation, have fundamental roles in regulating several cellular processes ranging from cancer to infectious diseases^14,15^. In particular, histone acetylation, which usually occurs at lysine residues at the N-terminus of histone proteins, is generally associated with transcriptional activation promoting euchromatin conformation, while histone deacetylation is generally associated with transcriptional repression by heterochromatin conformation^16,17^. The acetylation of lysine residues is regulated by lysine acetyltransferases (KATs) and by non- enzymatic reactions^18^. The opposite reaction is controlled by lysine deacetylases, or KDACs, which remove an acetyl group from lysine residues^19,20^. The classical KDACs are Zn^2+^- dependent while sirtuins are NAD^+^-dependent deacetylases, and together with KATs maintain a balanced acetylation status of proteins.

Recently, acetylation of histone and non-histone proteins has been investigated for its role in the regulation of gene expression, developmental processes, metabolism, stress resistance, pathogenesis, and virulence in some filamentous fungi^21–39^. Genetic knockout of sirtuin HstD/AoHst4 in *A. oryzae* interfered with fungal growth, sporulation, stress responses, and the production of secondary metabolites^28,29^. *A. nidulans* HstD/AoHst4 orthologue (AN1226) deletion resulted in decreased mycelial autolysis, conidiophore development, sterigmatocystin biosynthesis, and production of extracellular hydrolases^38^. Moreover, sirtuins were correlated with virulence, growth, and SMs biosynthesis in *A. flavus*^39^.

Although classical KDACs have been involved in different biological processes in *A. fumigatus* and virulence^23,35^, the biological role of sirtuins remains unexplored. This study aims to fill this knowledge gap by examining the role of sirtuins in *A. fumigatus* and their impact on key biological processes and virulence. *In silico* analysis indicated that *A. fumigatus* has six genes encoding putative sirtuins and their deacetylase activity was confirmed in four of them by *in vitro* assays using recombinant proteins. By combining gene knockout and extensive phenotyping of single mutant strains (ΔAfSirA, ΔAfSirB, ΔAfSirC, ΔAfSirD, ΔAfSirE, and ΔAfHstA), together with transcriptomic profiling, we demonstrate that despite sirtuins are not essential for *A. fumigatus* survival, they are involved in several biological processes. Our findings indicate their involvement in virulence, thermotolerance, SMs production, and cell wall biosynthesis. Additionally, acetylome data obtained from a single mutant strain (ΔAfSirE) and the null sirtuins strain (SIRTKO) provided valuable insights into the direct or indirect regulation of acetylation balance by sirtuins. Significant changes in the acetylation status of numerous proteins were observed, including histones and genetic determinants of virulence. These findings demonstrate the importance of sirtuins in *A. fumigatus* biology and pathogenesis. Understanding the role of sirtuins in this human pathogenic fungus may pave the way for effectively developing novel therapeutic strategies to combat *A. fumigatus* infections, particularly in immunocompromised patients.

## RESULTS

### *A. fumigatus* harbors six sirtuin genes

Six putative genes encoding sirtuins were identified in *A. fumigatus* genome (FungiDB) using the SirA sequence from *A. nidulans* (AN10449) as a query. These six proteins have the characteristic sirtuin domain and are grouped into four sirtuin classes (Class Ia, Ib, Ic, and III), as well as *S. cerevisiae* orthologs **(Figure 1A and B)**. The analyses of amino acid identity of *A. fumigatus* sirtuins (full length and sirtuin domain) showed a degree of conservation ranging from 17 to 49% and 25 to 57% of identity considering the sirtuin domain relative to *S. cerevisiae* and human orthologs, respectively (**Figure 1C and Sup. Figure 1A**). Moreover, amino acid sequence analyses demonstrated the presence of *A. fumigatus* sirtuin orthologues in several *Aspergillus* species (human pathogenic or not) with a high degree of conservation **(Sup. Figure 1B and C)**.

**Figure 1.**
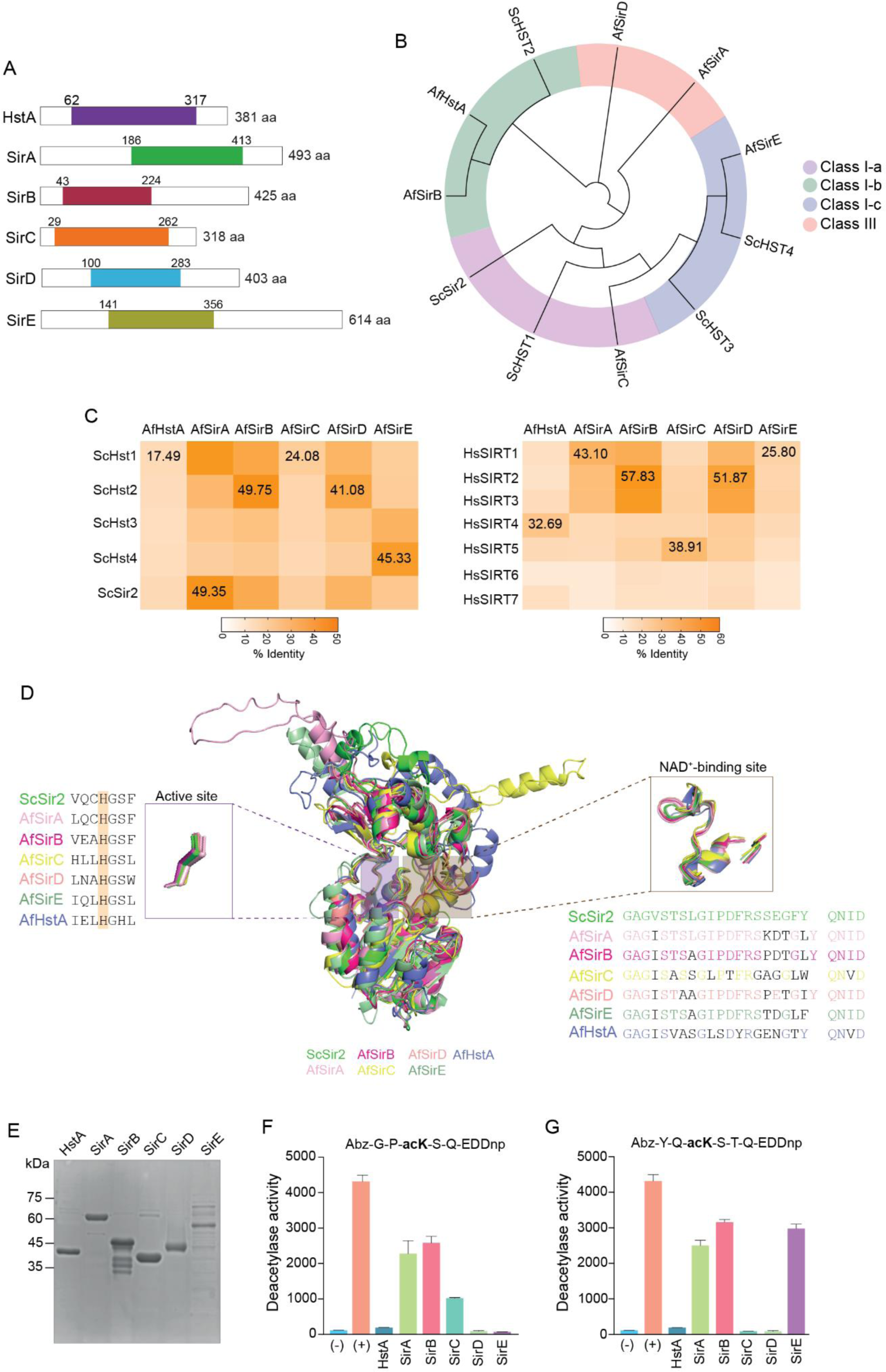
*A. fumigatus* sirtuins are active deacetylases. **A)** *In silico* analysis showed that *A. fumigatus* harbors six sirtuins that are schematically represented. The sirtuin catalytic domains are colored and flanked by distinct N- and C-terminal extensions. The numbers below each domain indicate the amino acid position for orientation. **B)** Phylogenetic tree of *A. fumigatus* (Af) and *S. cerevisiae* (Sc) sirtuins. For both species, the sirtuins are divided into Class Ia, Ib, II, and III. **C)** Identity (%) of full sirtuin sequences among *A. fumigatus*, *Saccharomyces cerevisiae* and human. **D)** Tridimensional structural analyses of predicted *A. fumigatus* sirtuin domains compared to *S. cerevisiae* homologs indicating the catalytic site with conserved histidine and NAD+ binding region. **E)** SDS-PAGE gel of purified *A. fumigatus* recombinant sirtuins used in the *in vitro* deacetylation assays. Expected sizes for each protein: HstA (41.5 kDa); SirA (54.6 kDA); SirB (46.9 kDa); SirC (34.6 kDa); SirD (45.1 kDa); SirE (67.1 kDa). **F-G)** *In vitro* deacetylase activity of *A. fumigatus* recombinant sirtuins using two acetylated peptides as substrate. HstA and SirD were the only enzymes that did not present deacetylase activity. The positive control used in the assays was the *Trypanosoma cruzi* Sir2rp1.

To gain insight into the characterization of *A. fumigatus* sirtuins, 3D predicted structural models for all enzymes were generated using the AlphaFold tool. All predicted models presented high confidence **(Sup. Figure 2)** and were used for comparative structural analyses with *S. cerevisiae* Sir2. The sirtuin domain of all *A. fumigatus* proteins has a high structural conservation degree compared to *S. cerevisiae* enzyme, including the key histidine involved in enzymatic activity and the residues of NAD^+^ binding site (**Figure 1D**).

To validate the deacetylase activity of *A. fumigatus* sirtuins, the six sirtuin-encoding genes were expressed in *E. coli* and used to test *in vitro* deacetylase activity (**Figure 1E**). Two acetylated peptides were designed to test the deacetylase activity of the *A. fumigatus* sirtuins, one based on the histone H3K56 acetylated site from *A. nidulans* (YQacKSTQ)^38^ and another general peptide (GPacKSQ). Recombinant sirtuins showed *in vitro* deacetylase activity, except HstA and SirD **(Figure 1F-G)**. Interestingly, while SirA and SirB were active on both peptides, SirC was active only in the general peptide (**Figure 1F**), and SirE showed specificity for the histone H3-derived peptide (**Figure 1G**), demonstrating a substrate-dependent activity of *A. fumigatus* sirtuins.

### Sirtuin-encoding genes are not essential to *A. fumigatus* but are involved in cell wall integrity, thermotolerance, and protease secretion

Six *A. fumigatus* single knockout strains (ΔAfSirA, ΔAfSirB, ΔAfSirC, ΔAfSirD, ΔAfSirE, and ΔAfHstA) in addition to a strain carrying the deletion of the six sirtuins (SIRTKO) were constructed using the CRISPR-Cas9 system available for *Aspergilli*^40,41^ . The deletions were confirmed by PCR and Southern blot analyses (**Sup. Figure 3 and Sup. Figure 4**). All the single mutants were viable under lab conditions; however, ΔAfSirE and SIRTKO displayed significant growth defects in yeast agar glucose (YAG) medium (**Figure 2A**). The influence of sirtuins on cell wall integrity was evaluated by exposing the mutant strains to cell wall stressors. ΔAfSirA, ΔAfSirB, ΔAfSirE, and SIRTKO strains showed sensitivity to cell wall stressors, such as Congo Red (CR), Calcofluor White (CFW), and caspofungin, compared to the WT strain (**Figure 2B-D**). Moreover, analyzing the monosaccharides in the cell wall revealed significant differences in the contents of glucosamine in ΔAfSirA, ΔAfSirB, and ΔAfHstA strains and galactose in ΔAfSirA, ΔAfSirB, ΔAfSirC, and ΔAfSirE mutants compared to the WT strain, but no significant differences in mannose content **(Figure 2E)**. In addition, all sirtuin mutants showed resistance to high temperatures **(Figure 2F)**. The ΔAfSirE and SIRTKO strains demonstrated greater sensitivity to caspofungin and voriconazole incorporated into a solid medium. In addition, complete inhibition of macroscopic growth was observed at 2 μg/mL of caspofungin, rather than partial inhibition defined as the Minimal Effective Concentration (MEC) for caspofungin (**Sup. Figure 5A and B**).

**Figure 2.**
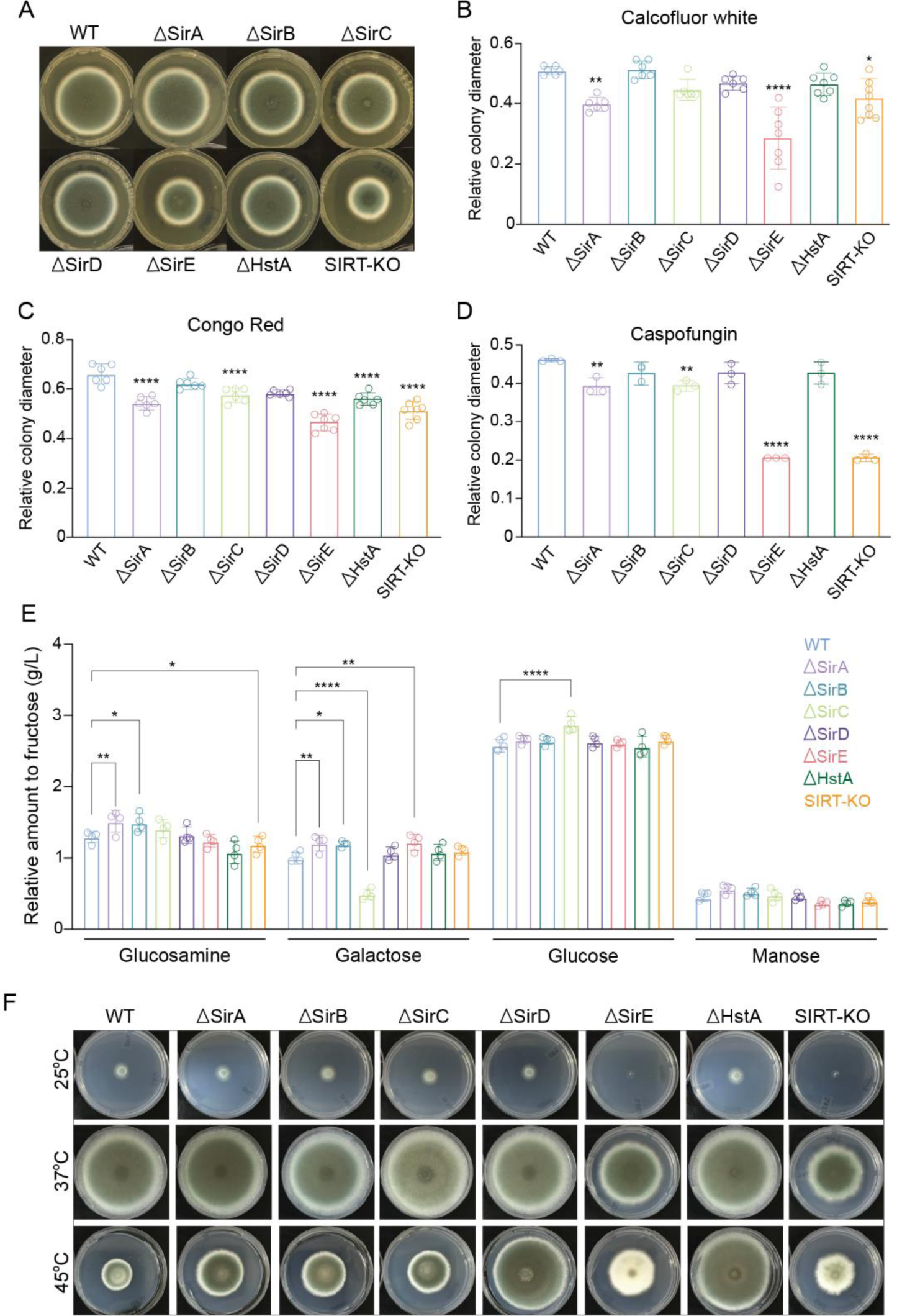
Phenotypic characterization of *A. fumigatus* sirtuin mutants. **A)** *A. fumigatu*s mutants inoculated on YAG medium for 120 h at 37°C. **B)** Relative quantification of mutant strains growth based on colony diameter in the presence of calcofluor white (30ug/mL, n=6), **C)** congo red (50 µg/mL, n=6), and **D)** caspofungin (1ug/mL, n=3). **E)** Cell wall polysaccharides composition of WT and sirtuin mutants mycelia after acid (H_2_SO_4_) treatment and quantification of monosaccharides (glucose, mannose, glucosamine, and galactose) by high-performance ionic chromatography (n=3) relative to fucose (internal control). **F)** *A. fumigatus* sirtuin mutants grown on Glucose Minimal Media (GMM) for 120 h at different temperatures. Significant differences were observed by using two-way ANOVA. *p < 0.05, **p < 0.002, ***p < 0.001 and ****p < 0.0001 indicates significant differences from comparisons to the WT strain.

### *A. fumigatus* sirtuins are involved in virulence

To access the virulence profile of sirtuin mutant strains, we reinserted the native *pyrG* gene in each mutant strain, due to the avirulent profile of uracil/uridine auxotroph strains^42^. *G. mellonella* larvae were infected with the conidia of each mutant strain, and the survival was evaluated. Interestingly, the ΔAfSirA, ΔAfSirB, ΔAfSirC and ΔAfHstA strains were hypervirulent and the ΔAfSirE and SIRTKO strains were less virulent when compared to the WT strain (**Figure 3A**). These results indicate that sirtuins have an impact on *A. fumigatus* virulence in a non-vertebrate model. Due to the attenuated virulence of ΔAfSirE strain, an additional assay was carried out using a murine model (**Figure 3B**). To do that, we constructed an *AfsirE* complemented strain (ΔsirE::sirE) and a strain carrying an inactive version of SirE, substituting a histidine (H) at position 260 with tyrosine (Y) (ΔAfSirE^H260Y^) by site-directed mutagenesis (**Figure 1B**). ΔAfSirE, ΔAfSirE^H260Y^, and SIRTKO strains were avirulent in the neutropenic mouse model, while virulence of the WT strain was partially recovered in the complemented strain ΔsirE::sirE. The inactivation of AfSirE^H260Y^ was confirmed by testing the *in vitro* deacetylation activity of the heterologous protein compared to the native protein **(Sup. Figure 6)**. In line with mortality, the mutant strains SIRTKO, ΔAfSirE, and ΔAfSirE^H260Y^ displayed a less accentuated body weight loss (**Figure 3C**), and less pronounced clinical signs in comparison to the WT strain (**Figure 3D**).

**Figure 3.**
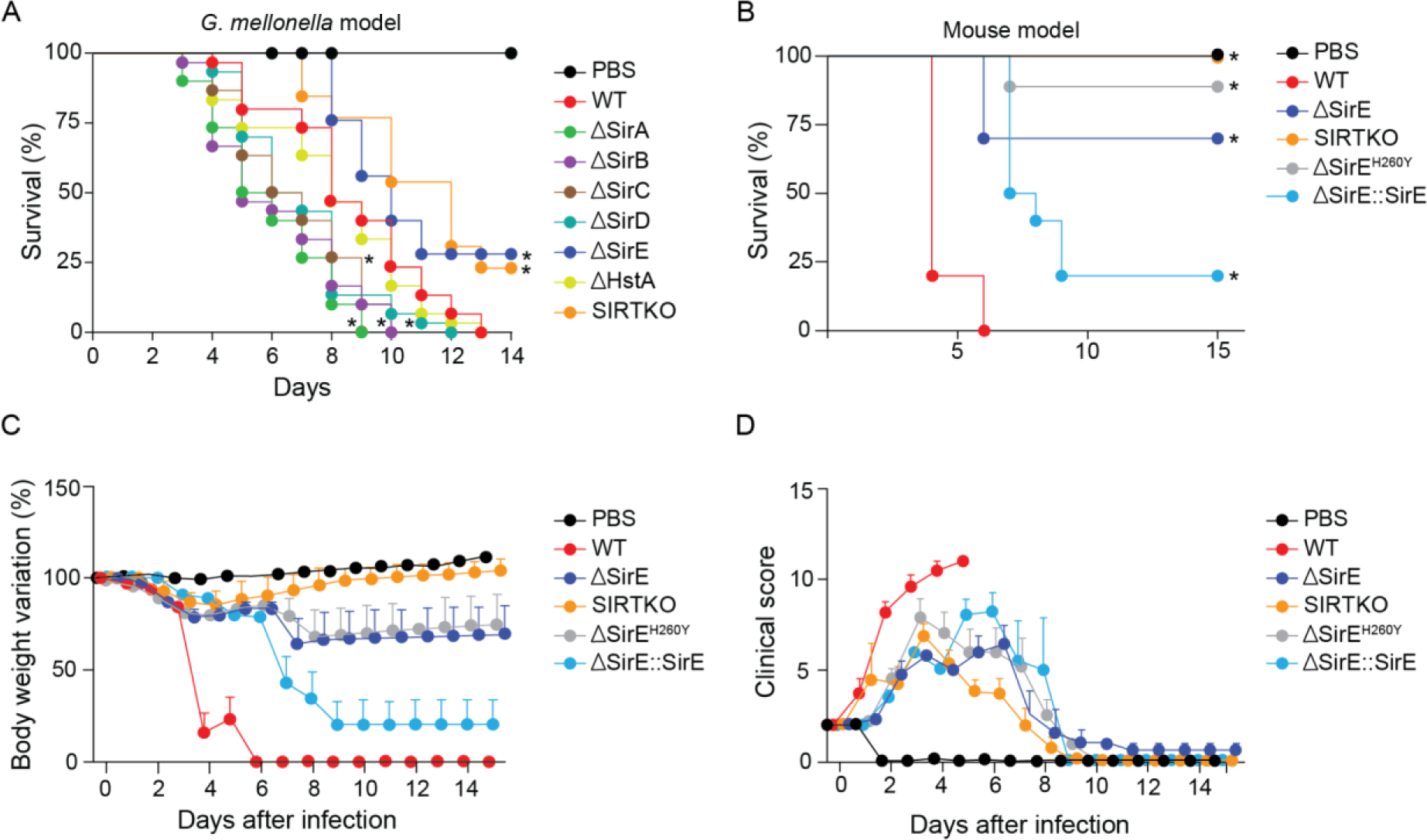
Virulence assay. **A)** Survival curve of *A. fumigatus* sirtuin mutant strains using *Galleria mellonella* model. The graph represents the average of three independent replicates using ten larvae for each experiment. **B)** Survival curve of WT, ΔAfSirE, ΔAfSirE^H-Y^ and ΔsirE::sirE complemented strains in female BALB/c mice. **C)** Body weight variation of infected animals with the different *A. fumigatus* strains. **D)** Clinical Score of animals during the course of infection. Significant differences were observed by using a two-way ANOVA *p < 0.05.

### Sirtuins regulate secondary metabolite production

The metabolome of sirtuin knockout strains was analyzed to understand the impact of sirtuins on the regulation of SMs in *A. fumigatus*. Principal component analysis (PCA) was applied to examine data acquisition reproducibility, batch carry-over effects, and chemical-based grouping tendencies (**Sup. Figure 7A**). After ensuring the clustering of QC samples near the origin of the coordinate system, which indicates good analytical method reproducibility, a PCA model was constructed with the samples of only the fungal strains (**Figure 4A**). The 2D score plot (**Sup. Figure 7B**) illustrates the clustering of WT, HstA and SirtA-D groups, indicating that a small metabolic variation resulted from the knockout of these genes. On the other hand, the metabolite profile of SIRTKO and SirtE was clearly distinct as observed across principal component 1. To identify SMs altered in SIRTKO and SirtE, structural annotation of chemical features was performed by either querying acquired MS/MS spectra against the Global Natural Products Social Molecular Networking (GNPS) public database or propagating annotations across molecular networks based on accurate mass and fragmentation similarity of well-established *A. fumigatus* SMs. Relative quantification based on the feature area of annotated SMs is represented on a heatmap **(Figure 4B and Sup. Figure 8)**. The data indicated that Pseurotin A, Brevinamide F, Pyripyroropene A, Deacetyl-Pyriopyropene and Bis(methylthio)gliotoxin were more abundant in both SIRTKO and ΔAfSirE strains, while Fumiquinazoline E and C, Trypostin B, Pyripiropene E-H, and Fumagillin were found in low abundance compared to the WT strain (**Figure 4B**). These data demonstrate that sirtuins affect the production of SMs in *A. fumigatus*.

**Figure 4.**
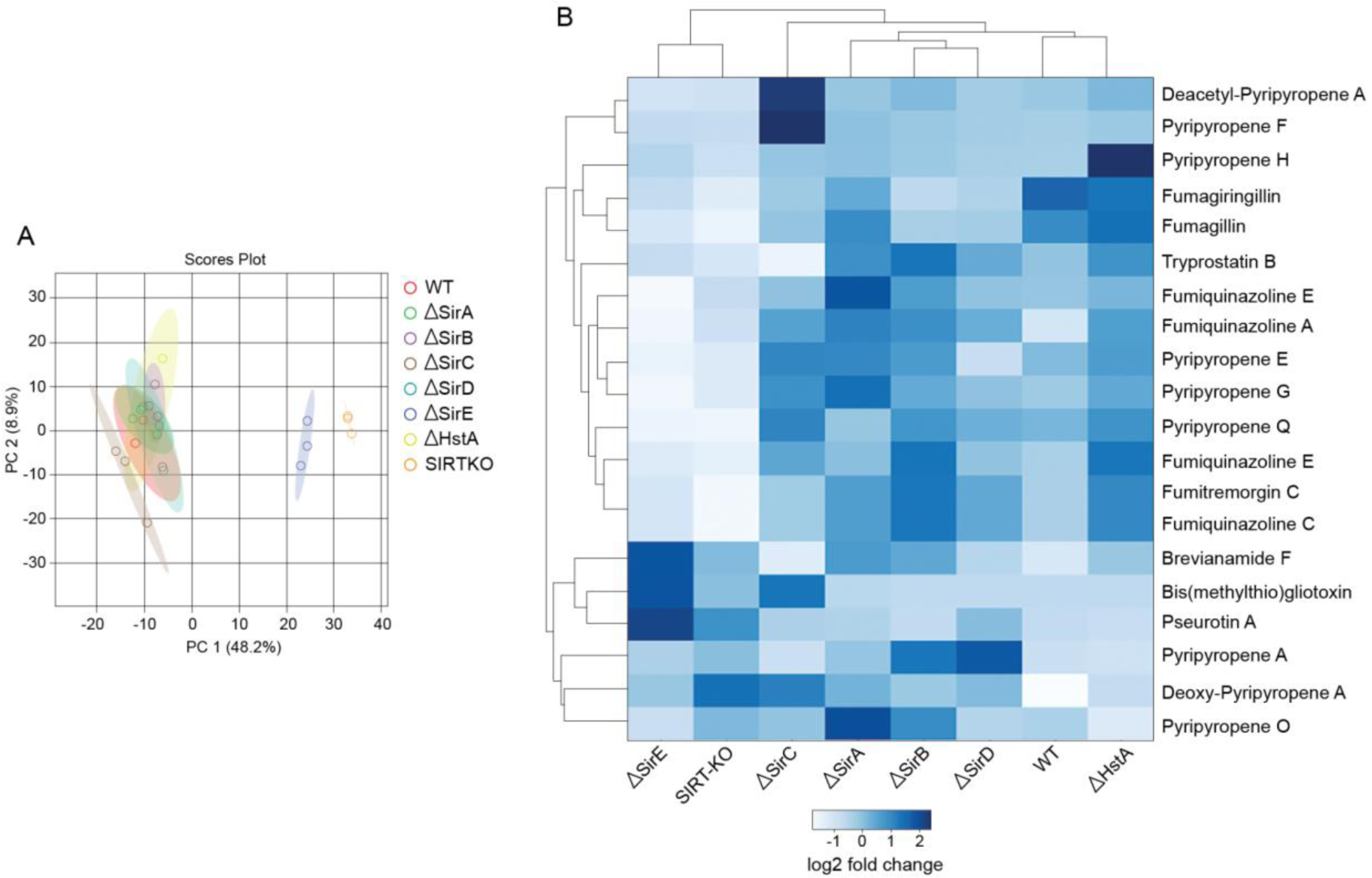
**A)** Principal component analysis (PCA) of the metabolic extracts of *A. fumigatus* sirtuin mutant strains. PCA model after removal of QC, analytical blanks and GMM media samples. **B)** Relative quantification of annotated secondary metabolites based on feature area. Data was acquired from an average of three biological replicates.

### Sirtuin deletion affects the acetylation abundance of proteins

Quantitative acetylome of the avirulent strains (ΔAfSirE and SIRTKO) was performed by enrichment analysis using the PTMScan® Acetyl-Lysine Motif [Ac-K] Kit (Cell Signaling Technology). The intracellular proteins of WT and mutant strains were extracted, then digested with trypsin, and either the total proteome and acetyl-lysine (Kac)-enriched fraction were analyzed by nLC-MS/MS.

A total of 1548 proteins, or 15.7% of the *A. fumigatus* proteome, were identified in the SIRTKO and WT strains, while 957 proteins were found in the Kac-enriched fraction, with 390 of those proteins containing 598 Kac sites (**Figure 5A**). Our findings showed that 72.7% of the acetylated proteins contained one Kac site, while 27.3% were acetylated at two or more lysine residues (**Figure 5B**). Moreover, 159 Kac sites were distributed among 118 acetylated proteins in the WT strain, while 318 acetylated proteins displayed 439 Kac sites in the SIRTKO strain. Moreover, 295 Kac sites were found differentially acetylated in 216 proteins (False Discovery Rate (FDR) < 0.5; **Figure 5C and Sup. Table 4**). Among them, we found different abundances in histone acetylation. Histone H3 was abundantly acetylated at positions H3K56, H3K9, H3K36, H3K27, and H3K14, while lower acetylation in histone H2B was detected at position H2BK14, H2BK7, H2BK19, H2BK24 in the SIRTKO strain. The set of differentially acetylated proteins was used as input for Gene Ontology (GO) enrichment analysis (**Figure 5D**), resulting in some overrepresented biological processes (BP) such as regulation of cellular component organization (GO:0016043), gene expression (GO:0010467), response to stress (GO:0033554), macromolecule metabolic process (GO:0019222), growth (GO:0040007), cellular localization (GO:0051641), and asexual reproduction (GO:0019954). The enriched cellular components (CC) range from the nucleus to the cell wall, whereas for molecular function (MF) there were enriched terms such as translation factor activity, RNA and histone binding, and N-acetyltransferase activity (**Figure 5D**). Seven genetic determinants of virulence were abundantly acetylated in the SIRTKO acetylome, these proteins are involved in metabolism, cell wall integrity, and signaling, such as the Bzip developmental regulator (Afu2g14680/flbB), putative mitogen-activated protein kinase kinase kinase - MAPKKK (Afu3g1108/bck1), UDP-galactopyranose mutase (Afu3g12690/glfA), cross-pathway control WD-repeat protein (Afu4g13170/ cpcB), GTP-Binding nuclear protein (Afu6g13300/Ran), and putative septin (Afu7g05370/aspB).

**Figure 5.**
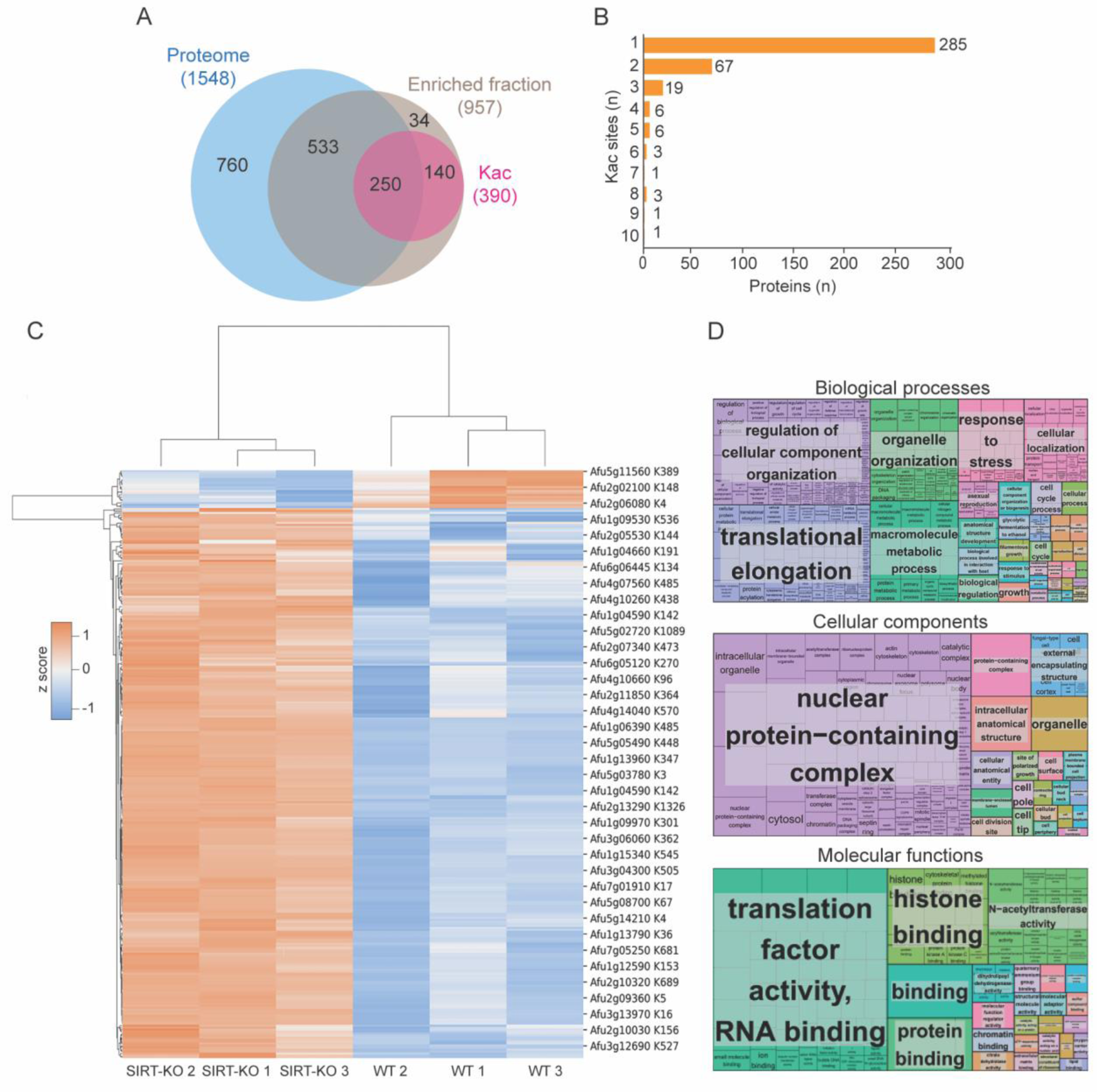
Acetylome analysis of *A. fumigatus* SIRTKO strain. **A)** Venn diagram of proteins found in the total proteome, enriched fraction, and lysine-acetylated proteins of WT and SIRTKO strains. **B)** Distribution of the number of acetylated sites (Kac) in the proteome. **C)** Heat map of Kac sites abundantly detected comparing the WT and SIRTKO strains. **D)** GO enrichment of the 216 proteins with differentially abundant Kac sites was performed using FungiDB and visualized with REVIGO to eliminate redundant terms (p-value cutoff> 0.05).

In addition to the profile of protein acetylation, 148 proteins were differentially expressed in the SIRTKO strain relative to the WT strain **(Sup. Table 3)**. These proteins are enriched for nucleotide metabolic process (GO:0009117), response to stress (GO:0006950), small molecule metabolic process (GO:0044281), and other biological processes (**Sup. Table 3**).

Around 1229 proteins were detected in the ΔAfSirE and WT proteomes, and 835 proteins were identified in the enriched fraction, in which 262 Kac sites were distributed among 170 proteins (**Figure 6A**). Approximately 71% of these acetylated proteins have only one acetylation site (**Figure 6B**). Moreover, 46 Kac sites were found differentially acetylated in 40 proteins of the ΔAfSirE strain (**Figure 6C**). GO terms enrichment analysis indicated that these proteins are involved with the regulation of defense response (GO:0031347), chromatin remodeling (GO:0006338), pyruvate metabolism (GO:0006090), as well as the GO terms cytosol (GO:0005829), DNA packing complex (GO:0044815), protein-containing complex (GO:0032991), cell wall (GO:0005618), binding (GO:0005488), and fatty acid synthase activity (GO:0004312) were also enriched (**Figure 6D**). Histones H3, H2A and H4 were abundantly acetylated (H3K56; H3K18; H3K23; H2AK6; H2AK4; H4K5) in the ΔAfSirE acetylome compared to the WT strain, while a lower acetylation abundance was found for histone H2B (H2BK7; H2BK14). In addition, 61 proteins were differentially expressed in the ΔAfSirE proteome, in which 42 hits were upregulated and 19 downregulated (**Sup. Table 3**). These differentially expressed proteins are enriched for pyrimidine deoxyribose metabolic and biosynthetic process, stress response, COPII-coated vesicle cargo loading, fatty acid metabolic process, and homocysteine biosynthesis (**Sup. Table 3**).

**Figure 6.**
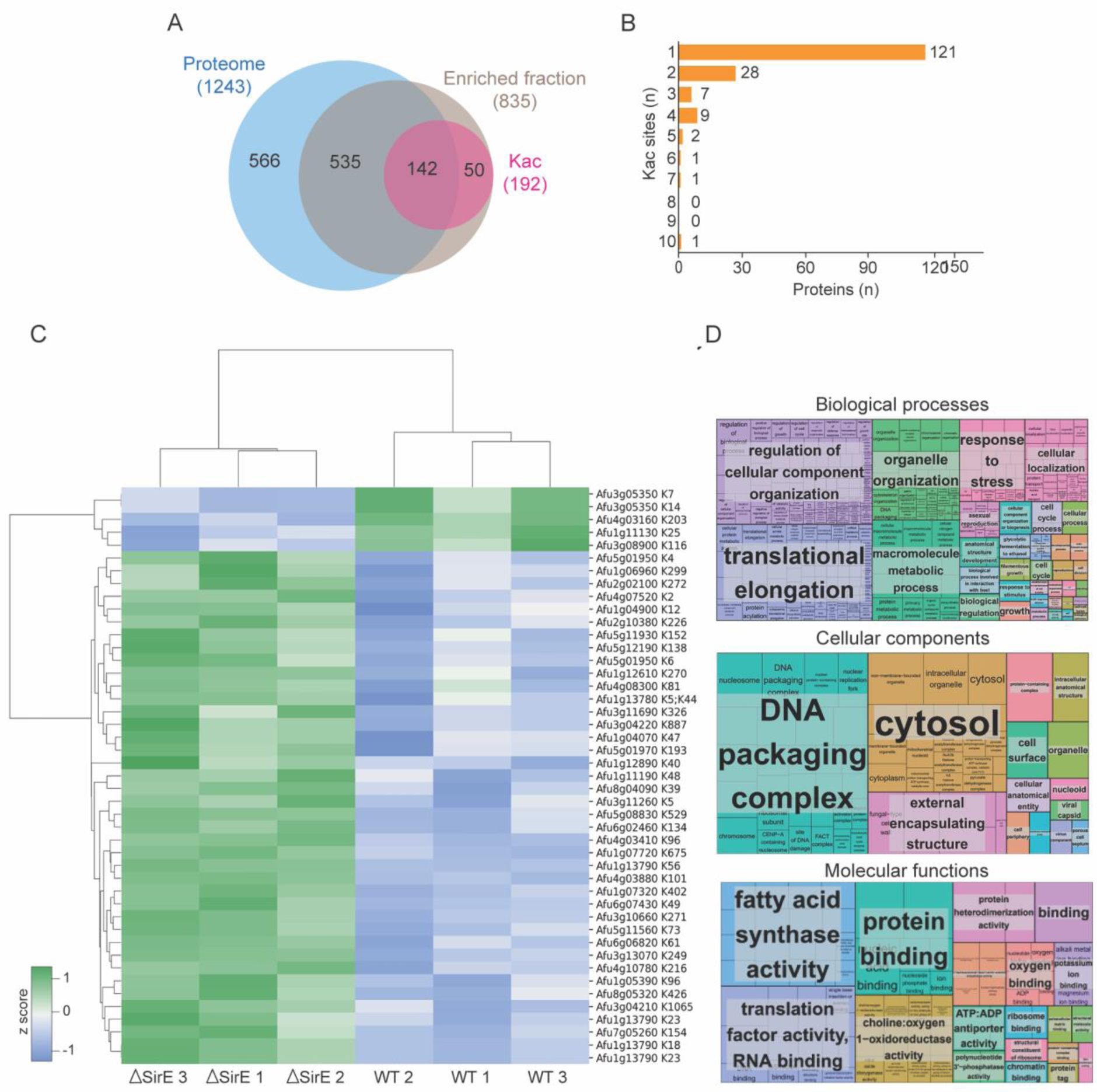
Acetylome analysis of *A. fumigatus* ΔAfSirE strain. **A)** Venn diagram of proteins found in the total proteome, enriched fraction and lysine-acetylated proteins of WT and ΔAfSirE strains. **B)** Distribution of the number of acetylated sites (Kac) in the proteome. **C)** Heat map representing the abundance of lysine-acetylated (Kac) proteins. **D)** GO enrichment of the 40 proteins with differentially abundant Kac sites was performed using FungiDB and visualized with REVIGO to eliminate redundant terms (p-value cutoff >0.05).

### Sirtuins regulate the transcription of genetic determinants of virulence

To understand whether sirtuins regulate gene expression, a comparative transcriptome analysis of WT, ΔAfSirE and SIRTKO strains cultured on GMM medium for 36 h was performed. The volcano plot and GO enrichment analysis of differentially expressed genes (DEGs) of *A. fumigatus* are represented in **Figure 7**. We identified 727 DEGs on the ΔAfSirE strain, of which 480 were upregulated and 247 downregulated compared to the WT strain (mean log 2-fold change ≥ 2 and FDR ≤ 1e^-3^) (**Figure 7B**). GO enrichment analysis indicated that among upregulated genes the most enriched biological process was associated with secondary metabolic processes (GO:0019748) (**Figure 7C and Sup. Table 5 and 6**). The downregulated genes were associated with transmembrane transport (GO:0072348), N-acetylglucosamine metabolic process (GO:0006044), glucosamine catabolic process (GO:0006043), and amino sugar catabolic process (GO:0046348) (**Figure 7D and Sup. Table 5 and 6**).

**Figure 7.**
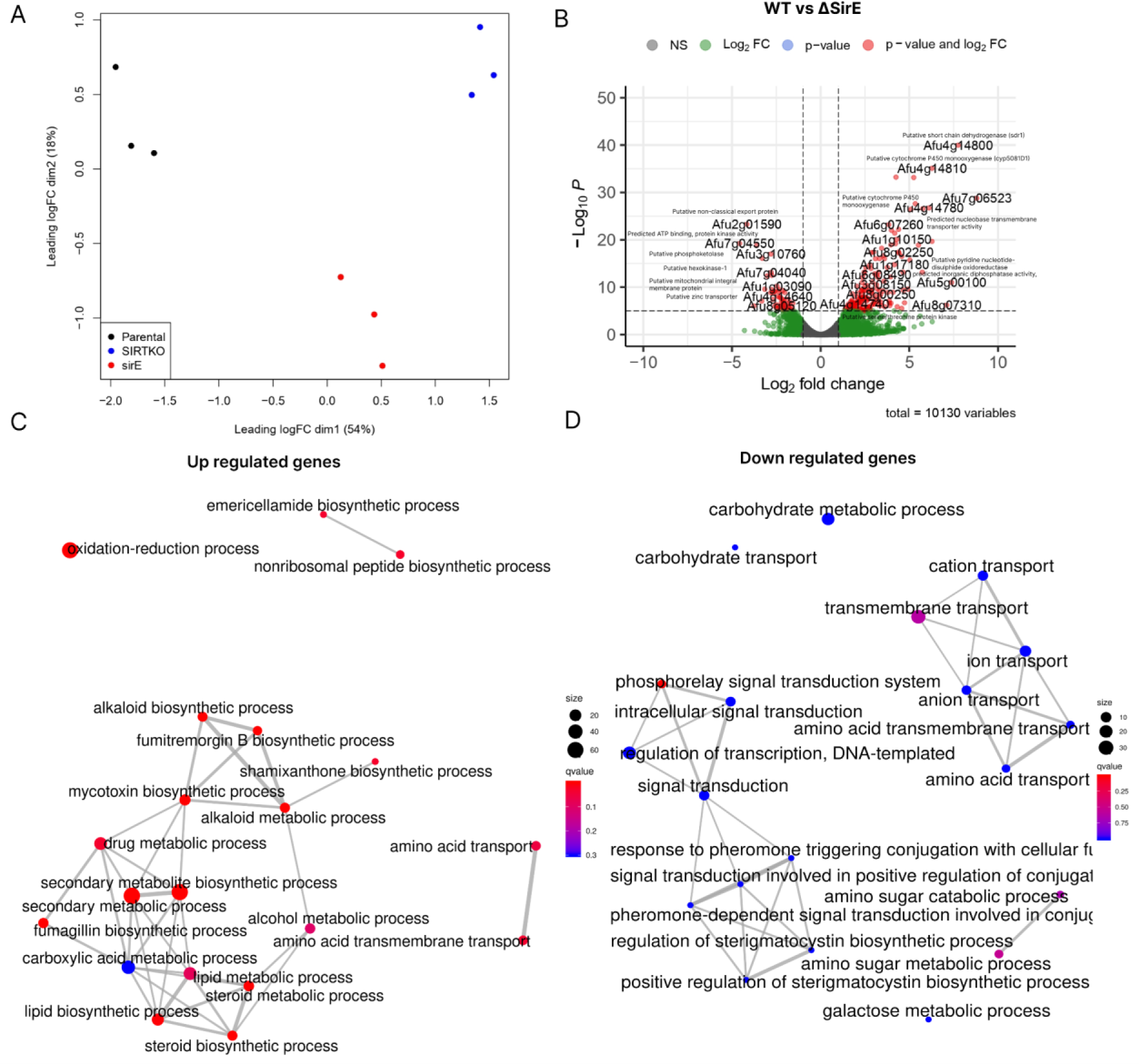
Transcriptome analysis of *A. fumigatus* Wt and ΔAfSirE strains. **A)** Multi-dimensional scaling (MDS) plot of RNA-seq datasets from WT, ΔAfSirE, and SIRTKO strain. **B)** Volcano plot of differentially expressed genes (|log2(FC)| > 1 and FDR < 1e^-5^). The most significantly expressed genes are labeled in red. **C)** Gene Ontology enrichment of upregulated and **D)** downregulated genes using the FungiExpresZ web platform for data visualization.

There are 36 genes potentially involved in *A. fumigatus* virulence upregulated in the ΔAfSirE strain **(Sup. Table 5)**. These genes are involved in the biosynthesis of SMs such as gliotoxin, fumagillin, brevianamide F, verruculogen, and fumitremorgin. Several putative cytochrome p450 monooxygenase enzymes (CYP450), which contribute to the production and diversity of various SMs^43^ were also upregulated in the ΔAfSirE strain. Among down- regulated genes in this strain, six are determinants of virulence, such as sensor histidine kinase/response regulator (Fos-1/TcsA; Afu6g10240), also found downregulated in response to itraconazole^44^, conidial hydrophobin (Afu5g09580), putative transporter (Afu3g12900), and the C6 transcription factor hasA (Afu3g12890) which controls the production of hexadehydroastechrome^45^.

On the SIRTKO strain, 1697 DEGs were identified, with 845 up- and 848 downregulated (**Figure 8B**). GO Enrichment analysis of upregulated genes showed DNA replication (GO:0006260), ketone metabolic process (GO:0042180), and SMs process (GO:0019748) as enriched biological processes (**Figure 8C and Sup. Table 5 and 6**), while the downregulated genes are involved in transmembrane transport (GO:0055085), N-acetylglucosamine catabolic process (GO:0006046), detoxification of inorganic compounds (GO:0061687), and regulation of stress-activated MAPK cascade (GO:0032872) (**Figure 8D and Sup. Table 4 and 5**).

**Figure 8.**
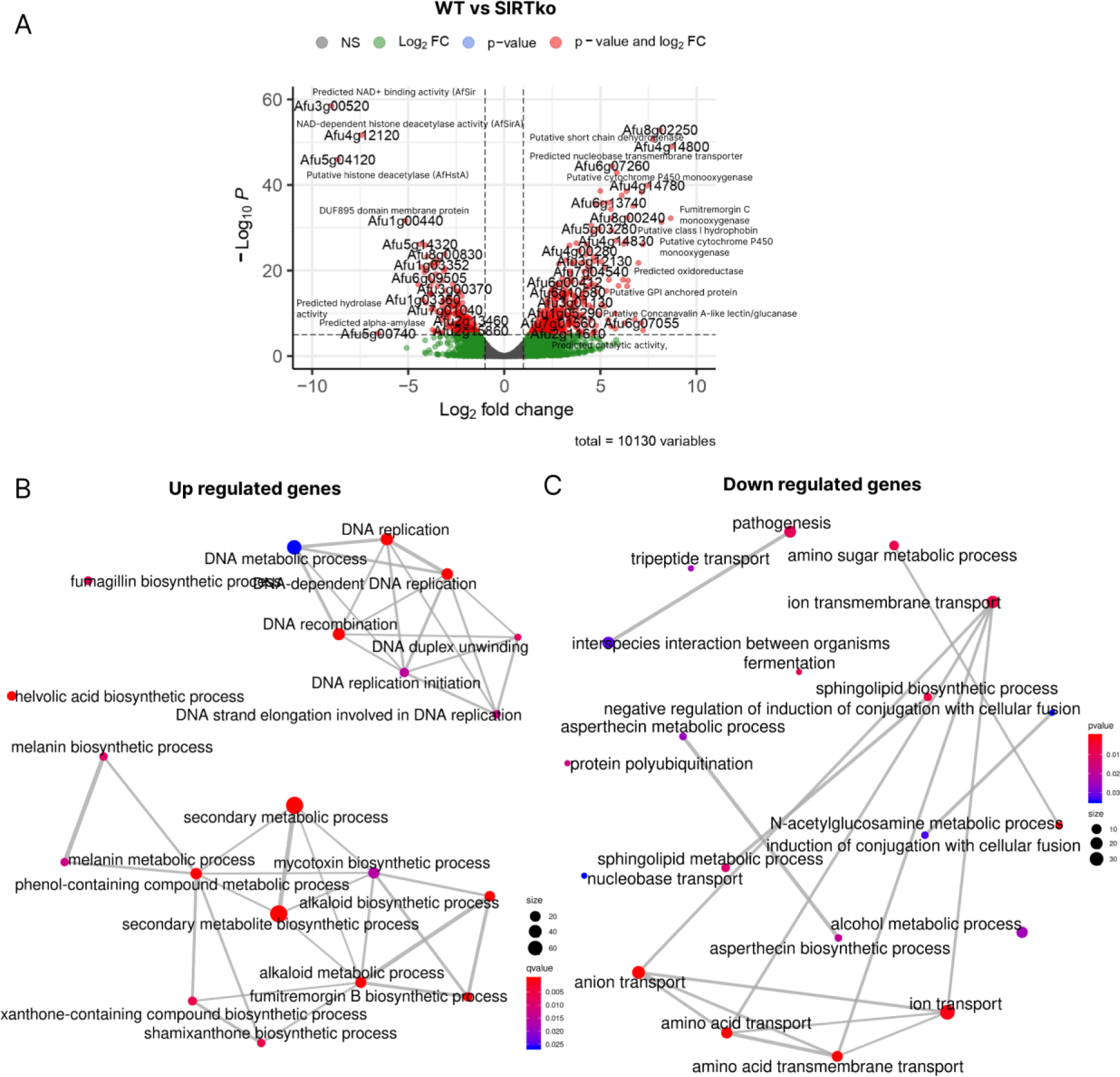
Transcriptome analysis of *A. fumigatus* WT and SIRTKO strains. **A)** Volcano plots of differentially expressed genes (|log2(FC)| > 1 and FDR < 1e^-5^). The most significantly expressed genes are labeled in red. **C)** Gene Ontology Enrichment for the biological function of upregulated and **D)** downregulated genes using the FungiExpresZ web platform for data visualization.

Among the DEGs in the SIRTKO strain, 36 and 20 genetic determinants of virulence were up and down-regulated, respectively. There are several genes involved in SMs biosynthesis such as heptaketide hydrolyase (*ayg1*/Afu2g17550), putative fatty acid oxygenase (Afu3g12120), conidial pigment polyketide synthase (*alb1*/Afu2g17600), as well as putative acetyltransferase with a predicted role in iron metabolism (Afu3g03650) upregulated in the SIRTKO strain. Among the 20 down-regulated genes, there are transcription factors involved in the regulation of morphogenesis, gliotoxin production, and virulence (Afu6g02690/*mtfA*), putative chitin synthase (Afu8g05630/*chsF*), putative regulator of adherence, host cell interactions and virulence (Afu2g13260/*medA*), and putative mitogen-activated protein kinase (Afu5g09100/*mpkC*).

In summary, taken together the transcriptome and acetylome analysis, and the in vitro deacetylase activity, our results suggests that sirtuin loss-of-function provided higher acetylation of histones inducing the upregulation of a large set of genes.

## DISCUSSION

Protein acetylation has been correlated to fungal growth, metabolism, pathogenesis, and SMs production^46–48^. The balance of acetylation and deacetylation of lysine residues can alter protein conformation and function, which can modulate several biological processes by influencing transcription regulation, enzyme activity, cell signaling, and protein-protein interactions^17^. Recent studies have suggested that sirtuins may play a role in virulence in bacteria and fungi. For example, sirtuins can modulate the expression of genes involved in virulence and pathogenesis in *Staphylococcus aureus*^49^ and *C. albicans*^50^.

In this study, we demonstrate the activity of four of six *A. fumigatus* sirtuins and the role of sirtuins in this pathogenic fungus by combining phenotypic characterization of six sirtuin knockout strains (ΔAfSirA-E and ΔAfHstA) and a strain displaying deletion of the six sirtuins (SIRTKO). SirE showed activity only on the histone H3-derived peptide, SirC was active only in the general peptide, and SirA and SirB were active on both peptides. All mutant strains constructed are viable under laboratory conditions, indicating that sirtuins are not essential genes. However, our results strongly suggest that sirtuins play roles in cell wall integrity, protease secretion, SMs production, and virulence. Notably, *A. fumigatus* ΔAfSirE and SIRTKO strains displayed a significant defect in radial growth, susceptibility to cell wall stressors, and attenuated virulence in both *G. mellonella* and mouse infection models. Due to the avirulent behavior of mutant strains, the profiles of both protein acetylation and gene expression were investigated. We found eight genetic determinants of virulence that could be sirtuin targets and hundreds of genes related to virulence that are differentially expressed. Among up- regulated genes, ketone metabolic process and secondary metabolism biosynthesis are enriched, while N-acetylglucosamine catabolic process and transmembrane transport were enriched among the down-regulated genes.

The virulence of *A. fumigatus* is attributed to multiple factors^51^. The components of cell wall, composed of layers of galactomannan, galactosaminogalactan, chitin, and beta-glucans^52,53^ contribute to the structural integrity of the cell wall and also play important roles in interactions with the host immune system^7^. Genes involved in cell wall biosynthesis are considered virulence factors and are frequently used as drug targets for treating fungal infections^54,55^. All *A. fumigatus* sirtuin mutants constructed displayed some sensitivity to one of the cell wall stressors tested, as well as differences in mycelial carbohydrate contents. In our transcriptome analysis, galactose biosynthesis and N-acetylglucosamine catabolic processes are enriched in the ΔAfSirE and SIRTKO strains considering the downregulated genes, corroborating the phenotyping of these strains. Interestingly, UDP-galactopyranose mutase (*glfA)*^56,57^, an important virulence factor and a potential target for new antifungal compounds, was found abundantly acetylated in SIRTKO Kac-enriched proteome, suggesting a possible regulation by sirtuins.

Filamentous fungi can produce a wide variety of SMs that contribute to their survival, fitness, pathogenicity, and virulence^11,58^. Histone acetylation status is an important regulator of chromatin structure, which impacts the expression of biosynthetic gene clusters in filamentous fungi^59,60^. Sirtuins have been reported as regulators of SM production in *Aspergillus* spp.^29,33,37,38,61,62^. The acetylome of *A. fumigatus* WT and SIRTKO strains demonstrate that the deletion of sirtuins promotes histone hyperacetylation and, therefore, could activate transcription of SMs gene clusters resulting in a different metabolome profile, which was already seen in other fungi^63^. Our transcriptome data also reveal hundreds of differentially expressed genes related to secondary metabolism, which corroborates the difference in the abundance of secondary metabolites observed in the mutant strains’ metabolome. Moreover, genes involved in various metabolic pathways and biological processes important to the pathogenicity of *A. fumigatus*, such as SM biosynthesis, cell wall integrity pathway, cell signaling, integral component of membrane, fatty acid metabolism, and pyruvate metabolism were differentially expressed in the ΔAfSirE and SIRTKO strains.

In conclusion, we show that sirtuins are involved in several biological processes, probably by maintaining protein acetylation and NAD^+^/NADH balance, which affect primary and secondary metabolism. Due to the loss of *sirE* in *A. fumigatus*, susceptibility to voriconazole and caspofungin increased. In addition, AfSirE has a low identity with human sirtuins, making it a potential drug target. Interestingly, nicotinamide (NAM) a classical sirtuin inhibitor, exhibited significant antifungal activity against *C. albicans* and had a synergistic interaction with amphotericin B against *C. albicans* as well as other *Candida* spp. and *Cryptococcus neoformans*^64,65^. Understanding the biology of these enzymes in *A. fumigatus* and how protein acetylation interferes with virulence may open perspectives for the development of new molecules targeting protein acetylation. Further *in vivo* studies will be necessary to assess the therapeutic potential of sirtuin inhibitors against the most common fungal pathogens.

## MATERIAL AND METHODS

### In silico analysis

The protein sequences of *A. fumigatus* sirtuins were retrieved from FungiDB database, using the following gene IDs: Afu5g04120 (AfHstA); Afu4g12120 (AfSirA); Afu2g05900 (AfSirB); Afu6g09210 (AfSirC); Afu3g00520 (AfSirD); Afu1g10540 (AfSirE). The *S. cerevisiae* and human sirtuin sequences were obtained from the UniProt database^66^. Then, using the respective sequences of the sirtuins from *A. fumigatus* and *S. cerevisiae*, the predicted 3D structures were generated using the artificial intelligence algorithm Alphafold Protein Structure Database developed by DeepMind and EMBL-EBI^67,68^. PyMOL software was used for structural analyses and to generate the images. The amino acid identity analyses and phylogenetic tree were done using the software Geneious Prime.

### Protein expression and purification

The pET28a-AfSirA, pET28a-AfSirB, pET28a-AfSirC vectors were transformed into *E. coli* BL21 (DE3) and the pET28a-AfHstA, pET28a-AfSirD and pET28a-AfSirE vectors were transformed into *E. coli* Arctic Express by heat shock and selected in solid LB medium in the presence of 50 µg/mL ampicillin. For AfHstA, AfSirA, AfSirB, AfSirC, AfSirD and AfSirE protein expression, 10 mL of an overnight grown culture was inoculated into 500 mL of liquid LB medium in the presence of 50 µg/mL kanamycin and maintained for 2h at 37°C and 180 RPM. After 2h of growth, 1 mM IPTG was added to the culture and it was conditioned overnight at 12°C and 180 RPM. The cultures obtained after induction of expression were centrifuged at 10.000 RPM for 15 minutes, resuspended with 10 mL of 20 mM Tris-HCl, pH 8.0, and centrifuged again. The bacterial precipitate was used in the protein purification steps.

Precipitates from the bacterial cultures obtained after induction were resuspended in lysis buffer (200 mM NaCl, 5% Glycerol, 5 mM 2-beta mercaptoethanol and 25 mM Hepes- NaOH, pH 7.5) and lysed using French Press apparatus. After centrifugation, the soluble fraction was used for the purification of heterologous proteins using Ni-NTA agarose resin (QIAGEN), previously equilibrated with equilibration buffer (200 mM NaCl, 5% Glycerol, 5 mM 2-beta mercaptoethanol, 25 mM imidazole, 25 mM HEPES-NaOH, pH 7.5), and incubated for 1h under stirring at 4°C with the soluble fraction of proteins previously obtained by French Press. After incubation, the resin was washed 4x with 10 mL of wash buffer (200 mM NaCl, 5% glycerol, 5 mM 2-beta mercaptoethanol, 50 mM imidazole, 25 mM HEPES-NaOH pH 7.5). Proteins were eluted from the Ni-NTA column with 1 mL of elution solution (200 mM NaCl, 5% glycerol, 5 mM 2-beta mercaptoethanol, 250 mM imidazole, and 25 mM HEPES-NaOH, pH 7.5). To confirm the quality of purified proteins, the samples were analyzed using acrylamide gel electrophoresis.

### Sirtuin deacetylase activity assay

Deacetylation assays were performed with purified recombinant AfHstA, AfSirA, AfSirB, AfSirC, AfSirD and AfSirE proteins following the protocol established by Moretti et al.^69,70^. Briefly, the substrate for enzyme activity consists of a peptide containing an acetylated lysine residue associated with a fluorescent group and an eraser group (Abz-Gly-Proacetyl-Lys-Ser-Gln-EDDnp), where Abz is ortho-aminobenzoic acid; and EDDnp is N-[2,4-dinitrophenylethylenediamine], respectively. In addition, another substrate was tested with all proteins and was synthesized based on the histone site H3K56ac, following the same conditions cited (Abz-Tyr-Gln-Proacetyl-Lys-Ser-Thr-Gln-EDDnp). The reaction is divided into two steps, in the first, corresponding to the deacetylation reaction; and the second, corresponding to the trypsinization of the peptide. Reactions were performed in 96-well dark plate, incubated for 4 hours at 37°C in 50 μL containing sirtuin activity buffer (25 mM Tris-HCl pH 8; 137 mM NaCl; 2.7 mM KCl; 1 mM MgCl2), 0.6 mM NAD^+^ (SigmaAldrich). After reaction time, a 12 mM nicotinamide solution in 100 mM NaCl and 50 mM Tris-HCl, pH 8, containing 0.6 mM trypsin (SigmaAldrich) was added to the reaction. After 30 minutes of incubation at 37°C, fluorescence (Ex 320 and Em 420 nm) was measured using SpectraMax M3 equipment (Molecular Devices, Sunnyvale, CA, USA).

### Strain and culture conditions

The *A. fumigatus* strains used in this study (Table 1) have been maintained in complete medium (YG; glucose 2% (w/w), 0.5% yeast extract (w/w), 1X trace elements) or glucose minimal medium glucose (GMM; glucose 1% (w / w), 1x high nitrate salt solution (1.4 M NaNO_3_, 0.13 M KCl, 0.042 M MgSO_4_ ·7H_2_O and 0.22 M KH_2_PO_4_) and 1x trace elements (7.2 mM ZnSO_4_ ·7H_2_0, 17.7 mM H_3_BO_3_, 2.52 mM MnCl_2_ ·4H_2_0, 2.72 mM FeSO_4_ ·7H_2_0, 0.95 mM CoCl_2_ ·5H_2_0, 0.7 mM CuSO_4_ ·5H_2_O, 0.21 mM Na_2_MoO_4_ ·4H_2_0 and 17.11 mM EDTA), pH 6.5) and (0.12% (w/v) uracil/uridine when required. For solid media, 1.5% agar (w/v) was added to YG or GMM, respectively. Plates were incubated at 37°C for 36 to 72 h.

**Table 1.** Strains used in this study.

| *A. fumigatus* strains | Genotype | Reference |
| --- | --- | --- |
| A1160- | *pyrG1*; $\Delta KU80$ | [Da Silva et al., 2006]<sup>69</sup> |
| $\Delta AfSirA$- | *pyrG1*; $\Delta KU80$; $\Delta Afu4g12120$ | This study |
| $\Delta AfSirB$- | *pyrG1*; $\Delta KU80$; $\Delta Afu2g05900$ | This study |
| $\Delta AfSirC$- | *pyrG1*; $\Delta KU80$; $\Delta Afu6g09210$ | This study |
| $\Delta AfSirD$- | *pyrG1*; $\Delta KU80$; $\Delta Afu3g00520$ | This study |
| $\Delta AfSirE$- | *pyrG1*; $\Delta KU80$; $\Delta Afu1g10540$ | This study |
| $\Delta AfHstA$- | *pyrG1*; $\Delta KU80$; $\Delta Afu5g04120$ | This study |
| SIRTKO- | *pyrG1*; $\Delta KU80$; $\Delta Afu4g12120$;<br>$\Delta Afu2g05900$; $\Delta Afu6g09210$;<br>$\Delta Afu3g00520$; $\Delta Afu1g10540$; $\Delta Afu5g04120$ | This study |
| $\Delta AfSirA^+$ | $\Delta KU80$; $\Delta Afu4g12120$; *pyrG1::pyrG* | This study |
| $\Delta AfSirB^+$ | $\Delta KU80$; $\Delta Afu2g05900$; *pyrG1::pyrG* | This study |
| $\Delta AfSirC^+$ | $\Delta KU80$; $\Delta Afu6g09210$; *pyrG1::pyrG* | This study |
| $\Delta AfSirD^+$ | $\Delta KU80$; $\Delta Afu3g00520$; *pyrG1::pyrG* | This study |
| $\Delta AfSirE^+$ | $\Delta KU80$; $\Delta Afu1g10540$; *pyrG1::pyrG* | This study |
| $\Delta AfHstA^+$ | $\Delta KU80$; $\Delta Afu5g04120$; *pyrG1::pyrG* | This study |
| SIRTKO<sup>+</sup> | $\Delta KU80$; $\Delta Afu4g12120$; $\Delta Afu2g05900$; $\Delta Afu6g09210$;<br>$\Delta Afu3g00520$; $\Delta Afu1g10540$; $\Delta Afu5g04120$;<br>*pyrG1::pyrG* | This study |
| $\Delta AfSirE^{H-Y}$ | $\Delta KU80$; *Afu1g10540* H113Y | This study |
| $\Delta AfSirE::sirE$ | $\Delta KU80$; $\Delta Afu1g10540::Afu1g10540$ | This study |

#### CRISPR-Cas9 vector construction

The construction of the CRISPR-Cas9 vector for a target gene was performed as previously described^40,41^. Briefly, small DNA fragments (biobricks) were amplified by touchdown PCR using a PfuX7 as polymerase^73,74,70–72^ and the vector pFC902 as template. The primers and repair oligonucleotides were designed using the *A. fumigatus* genomic sequence of each target (*Afu4g12120; Afu2g05900; Afu6g09210; Afu3g00520; Afu1g10540* and *Afu5g04120*) and are listed in **Sup. Table 1**. The PCR products were treated using DpnI (New England Biolabs) and purified using the Wizard SV Gel system and the PCR Clean-Up System (Promega).

The biobricks, which consist of a protospacer adjacent motif (PAM) sequence, were linked with a pFC330 (pyrG marker vector) by uracil-specific excision reagent (USER) fusion and USER cloning^75^. The mixture was used directly to transform thermo-competent *E. coli* DH5α cells. Transformants were plated on solid Luria Broth (LB) medium supplemented with 100 mg/ml ampicillin. Colonies were randomly selected for each construct and validated by colony PCR and PCR-product sequencing.

#### Transformation in *A. fumigatus* and diagnostic PCR

Protoplast preparation of *A. fumigatus* was performed using protocols previously described^73^. For transformation, 8 µg of CRISPR vector and 100 µM of repair oligonucleotide (90pb) were mixed with 100 µL protoplasts solution and 50 µL of PEG solution (25% (w/v)) in STC 50 (1.2 M sorbitol, 10 mM CaCl2 and 50 mM Tris-HCl pH 7.5). After 20 min on ice, 1 ml of PEG solution was added, and the mixture was incubated at room temperature for 20 min. Then, 3 mL STC 50 was added, and 1 ml of the suspension was poured onto a protoplast-recovery medium (supplemented with 1.2 M sorbitol). Plates were incubated at 37°C until the appearance of isolate colonies. Monosporic purification was performed three times to obtain genetically pure mutants. Diagnostic PCR was performed using the genomic DNA of WT and mutant strains. The primers used are provided in **Sup. Table 1**.

#### Southern blot

Genomic DNA was extracted from mutant strains using phenol-chloroform^74^. Briefly, approximately 80 µg of DNA was used for digestion with different restriction enzymes. The fragments were separated on 0.8% agarose gel and transferred by capillarity to the Amersham Hybond N+ membrane (GE Healthcare). DNA hybridization was performed with the Amersham AlkPhos Direct Hybridization Buffer, being detected using a multifunctional imaging system Typhoon FLA 9500 (GE Healthcare). Primers used for probe sequence are in Table S2.

#### Phenotype characterization

The sirtuin mutants designed by CRISPR Cas9 were grown for 120 h at 37°C in YG agar with cell wall stressors, such as Calcofluor White (CFW) and Congo Red (CR). The mutant strains were also cultivated in different temperatures (25°C, 37°C and 45°C) in GMM agar and subjected to antifungal susceptibility tests described below. The graphs of relative radial growth (treated/control) measurements were constructed by GraphPad Prism software. Significant statistical analysis was set using the Dunnett or ANOVA test and p-value < 0.05.

#### Cell wall polysaccharides extraction and sugar quantification

All sirtuin mutant strains were incubated in MM for 36h at 37°C. After incubation, the mycelia were washed with distilled water and lyophilized. Fungal cell wall polysaccharides were extracted from 10 mg dry-frozen mycelia as described previously^75^. Briefly, 10 µL of extracted samples and standard sugars were subsequently analyzed by high-performance liquid chromatography (HPLC) on a Dionex Bio-LC system equipped with a CarboPac PA1 anion-exchange column and a CarboPac PA guard column by an isocratic elution of NaOH 16 mM. Fucose was added as internal control.

#### Minimal Inhibitory Concentration (MIC)

The MIC of sirtuin mutant strains was determined following the Clinical and Laboratory Standards Institute M38-A2 guidelines^76^. The susceptibility to azoles was compared to the reference strain (VRZ range 0.25-1.0 µg/mL: ITZ range 0.25- 2.0 µg/mL and FLZ not susceptible). The MIC scores were obtained visually with the inoculums of 4.0 × 10^4^ conidia/mL after incubation at 37°C for 48 h. The MIC was defined as the lowest concentration resulting in 100% growth inhibition compared to the drug-free control. The Minimal Effective Concentration (MEC) of caspofungin was performed with GMM liquid media.

#### Virulence assay in *Galleria mellonella* model

*G. mellonella* survival was assayed as previously described^77,78^. *G. mellonella* larvae were kept in sterile glass flasks with modified lids, with a hole in the center covered with an ultra-fine stainless steel wire mesh for better ventilation. Ten healthy *G. mellonella* larvae of similar weight (approximately 330 mg) were used in each group. Each larva was inoculated with 10 µL of conidia from the WT strain and the mutant strains of *A. fumigatus* on the last left proleg, directly on hemocoel. PBS was used as a control. The larvae were kept at 37 °C in the dark and the mortality rate of larvae was monitored daily for 15 days; the larvae that did not present movement after touch stimulation were considered dead. Significant differences were observed by using a two-way ANOVA *p < 0.05.

#### Virulence assay in immunosuppression of female BALB/c mice

Virulence assays were done in immunosuppressed female BALB/c mice. Wild-type BALB/c female mice, aged 6–8 weeks, were kept in the Animal Facility of the Laboratory of Molecular Biology of the School of Pharmaceutical Sciences of Ribeirão Preto, University of São Paulo (FCFRP/USP), in a clean and silent environment, under normal conditions of humidity and temperature, and with a 12 h light and dark cycle. The mice were given food and water ad libitum throughout the experiments. The procedures adopted in this study were performed following the principles of ethics in animal research and were approved by the Committee on Ethics in the Use of Animals (CEUA) of the FCFRP/USP (Permit Number: 08.1.1277.53.6; Studies on the interaction of *Aspergillus fumigatus* with animals) from the University of São Paulo, Campus of Ribeirão Preto.

Mice were immunosuppressed with cyclophosphamide (150 mg per kg of body weight), which was administered intraperitoneally on days -4, -1, and 2 prior to and post-infection. Hydrocortisone acetate (200mg/ kg body weight) was injected subcutaneously on day -3. *A. fumigatus* strains were grown on minimum medium (MM) for 2 days prior to infection. Fresh conidia were harvested in PBS and filtered through a Miracloth (Calbiochem). Conidial suspensions were spun for 5 min at 3,000 x g, washed three times with PBS, counted using a hemocytometer, and resuspended at a concentration of 5.0 × 10^6^ conidia/mL. The viability of the administered inoculum was determined by incubating a serial dilution of the conidia on MM at 37°C. Mice were anesthetized by halothane inhalation and infected by intranasal instillation of 1.0 × 10^5^ conidia in 20 ml of PBS. As a negative control, a group of mice received PBS only. Mice were weighed every 24 h from the day of infection and visually inspected twice daily. The statistical significance of comparative survival values was calculated by the Prism statistical analysis package by using Log-rank (Mantel-Cox) Test and Gehan-Brestow-Wilcoxon tests.

#### Evaluation of Clinical Disease Score

Mice were evaluated daily for weight change and clinical signs of aspergillosis. Each signal (piloerection, dyspnea and ataxy) was scored 2, 3 or 4 points, respectively, and the sum of them, per mouse, corresponded to the daily clinical disease score. Mice that lost more than 5% and less than 10% of their weight in a 24 h period were assigned 1 point and those with a loss superior to 10% received 2 points which were added to the daily score. In the absence of weight reduction or other aspergillosis signs, the score was zero. The quantification of the mice score was performed via a visual evaluation of the signs, in each mouse separately, daily, usually by at least two blinded examiners. Those mice that reached humane endpoints were euthanized to minimize mice suffering, and their cause of death was considered as fungal infection. Statistical analyses were performed using Graphpad Prism® 6 software. For all variables, normal distribution and homogeneous variance were tested. When the distribution was considered normal and with homogeneous variance, the parametric ANOVA test with Bonferroni’s post-test was used for three or more groups. Results were expressed as mean ± SEM (standard error of the mean). The differences observed were considered significant when p was < 0.05.

#### RNA extraction and RNA-Seq Data analyses

10^6^ conidia/mL of WT, AfSirE and SIRTKO strains were cultured in 50 mL of MMG media at 37 °C for 36h. The mycelia were collected and frozen in liquid nitrogen and ground into a powder. The RNA extraction was performed using Trizol followed by Direct-zol RNA Miniprep Kits (Zymo Research). The RNA quality was measured in a bioanalyzer and samples with RIN > 7.5 were sent to sequence.

#### Quality control and alignment against the reference genome

RNA paired-end sequencing quality control was assessed through FastQC (www.bioinformatics.babraham.ac.uk/projects/fastqc) and multiQC^79^. An average of 45 million reads were sequenced per sample. Both adapters and low-quality bases (QV < 20) were trimmed from reads’ extremities using Trimmomatic^80^ with a minimum read length of 30 bp. All libraries were mapped against the “FungiDB-56_AfumigatusAf293_Genome.fasta” reference genome (retrieved from fungidb.org) through STAR aligner^81^ with default parameters. STAR-generated sorted BAM output files were used for assigning read counts to gene features with featureCounts^82^ with the following parameters: -s2 -p -M -O -fraction. We relied on the “FungiDB-56_AfumigatusAf293.gff” annotation file also downloaded from fungidb.org.

#### Differential expression

Read counts’ table generated by featureCounts was then used as input for differential expression (DE) analyzes relying on the EdgeR classic exact test (0.01 FDR threshold)^83^. DE pairwise comparisons were performed as WT-vs-ΔSirE and WT-vs-SIRTKO.

The same read count table was also submitted to a multi-dimensional scaling (MDS) analysis using the *plotMDS* function from the EdgeR package. EnhancedVolcano (https://bioconductor.org/packages/release/bioc/html/EnhancedVolcano.html) was employed for an overall DE visualization through volcano plots. All tools described in this paragraph were run under the R environment version 4.1.0.

#### Protein extraction

10^6^ conidia/mL of WT and SIRTKO *A. fumigatus* strains were cultivated in 50 mL of YPD (Yeast Peptone Dextrose) for 36h. Mycelia were frozen by liquid nitrogen and ground into a powder, followed by transfer to a 50mL falcon tube and sonication five times on ice using a high-intensity ultrasonic processor in 5mL of lysis buffer [8 M urea, 2 mM EDTA, 65 mM DTT, 30 mM nicotinamide, 3 μM trichostatin A, and 1mM PMSF]. The remaining debris was removed by centrifugation at 10,000 ×g at 4°C for 10 min. The protein was precipitated with cold 15% trichloroacetic acid for 2h at 4°C. After centrifugation at 10,000×g at 4°C for 10 min, the supernatant was discarded, and the precipitate was washed three times with cold acetone. The protein was redissolved in a buffer [8 M urea and 100 mM NH4CO3 (pH 8.0)], and protein concentration was determined using a Bradford method^84^. For trypsin digestion, 5 mg of protein were reduced with 5 mM DTT for 25 min at 56°C, alkylated with 14 mM iodoacetamide for 30 min at room temperature in the dark, and then diluted 1:5 with 50 mM NH_4_CO_3_. Trypsin was added at a 1:100 enzyme-to-protein mass ratio for overnight incubation at 37°C and the next day topped up with the same amount following an extra 4 h incubation at 37°C. Trypsin digestion was terminated by adding 0.1% TFA final concentration and desalted using Sep-Pak C18 500mg sorbent cartridges (Waters). Peptides dried by vacuum centrifugation, followed by LC-MS/MS analysis.

#### Affinity enrichment of lysine-acetylated peptides

PTMScan® Acetyl-Lysine Motif [Ac-K] Kit (CSL) was used to enrich 5 mg of protein per sample following the manufacturers and *Schilling, B. et al* (2019)^85^ protocols, followed by LC-MS/MS analysis. Briefly, peptide samples resuspended in IAP buffer were incubated with 100uL of antibody-bead conjugates specific for acetyl-lysine (Cell Signaling Technologies, PTMScan kits #13416). Immunoprecipitation proceeded overnight at 4°C with gentle mixing. Next, beads were washed twice with 1 mL ice- cold IAP buffer, and then thrice with 1 mL ice-cold IAP buffer. Bound peptides were eluted sequentially with 45 mL and then 55 mL of 0.15% TFA for 5 minutes each at RT. Eluted PTM peptides were then directly loaded onto C18 Stage Tips (Waters), desalted with 0.2% FA in water, and eluted with a solution containing 50% ACN, 49.8% water, and 0.2% FA. Eluted peptides were dried completely and stored at -80°C until further analysis.

#### LC-MS/MS

The peptide mixture from the biological replicates was analyzed by LTQ Velos Orbitrap mass spectrometer (Thermo Fisher Scientific, USA) coupled with liquid chromatography-tandem mass spectrometry using an EASYnLC system (Thermo Fisher Scientific).

#### SMs extraction and UHPLC-HRMS/MS analysis

The mutant strains (10^5^ spores) were cultivated in solid GMM media for 120 h at 37°C. After culturing, the total content from each Petri dish was cut into small pieces (2 x 2 cm) and metabolites were extracted with equal quantities of methanol (MeOH) by ultrasonic extraction at room temperature for 1 h. Supernatant was then vacuum-filtered, and the solvent was removed under reduced pressure. The same procedure was performed for the control culture medium. Final extracts were stored at -20°C. For sample preparation, 1 mL of HPLC grade MeOH was added to each extract and sonicated until complete dissolution. 500 µL of the obtained extracts were transferred to vials and diluted with HPLC grade MeOH to a total volume of 1 mL. Quality control (QC) samples were prepared with 55 µL of each sample and diluted to a final volume of 1 mL with HPLC grade MeOH. UHPLC-HRMS/MS positive mode analysis was performed in a Thermo Scientific QExactive Hybrid Quadrupole-Orbitrap Mass Spectrometer. As a stationary phase, a Thermo Scientific column Accucore C18 2.6 µm (2.1 mm × 100 mm × 1.7 µm) was used. Mobile phase was 0.1% formic acid (A) and acetonitrile (B). Eluent profile (A/B %): 95/5 up to 2/98 within 10 min, maintaining 2/98 for 5 min, down to 95/5 within 1.2 min and maintaining for 8.8 min. Total run time was 25 min for each run and flow rate of 0.3 mL.min^-1^. The injection volume was 6 µL. MS spectra were acquired with *m/*z ranges from 100 to 1,500, with 70,000 mass resolution at 200 Da, 1 microscan, and an AGC target of 16 with a maximum ion injection time set to 100 ms. Ionization parameters: sheath gas flow rate (45), aux gas flow rate (10), sweep gas flow rate (2), spray voltage (3.5 kV), capillary temperature (250° C), S-lens RF level (50) e auxiliary gas heater temperature (400° C). MS^2^ spectra were collected using a normalized collision energy of 30 eV, and the 5 most intense precursors per cycle were selected and measured with 17,500 mass resolution at 200 Da, 1 microscan, and an AGC target of 1^5^ with a maximum ion injection time for MS/MS scans set to 50 ms.

##### UHPLC-HRM/MS data processing and Feature-Based Molecular Networking (FBMN)

Raw UHPLC-HRMS/MS data were converted into mzML format files in MSConvert with 64-bit binary encoding precision and peak peaking^86^. Feature detection was performed in MZmine2 (v.2.53)^87^. Raw data, converted mzML data, MZmine2 batch queue describing data processing steps and parameters, MZmine2 output and sample metadata files were deposited into the MassIVE repository (accession number: MSV000087365). Multivariate analyses were performed with the MetaboAnalyst tool (v.4.0)^88^. Briefly, the features list was filtered if their relative standard deviation were > 25% in QC samples, followed by normalization by sum and pareto scaling. Principal component analysis (PCA) was used to evaluate the robustness of data acquisition and to explore the differences in chemical space of each strain. For FBMN ^87^ the data were filtered by removing all MS/MS fragment ions within +/−17 Da of the precursor m/z. MS/MS spectra were window-filtered by choosing only the top 4 fragment ions in the +/−50 Da window throughout the spectrum. The precursor and fragment ion mass tolerance were both set to 0.02 Da. A molecular network was then created where edges were filtered to have a cosine score above 0.65 and more than 4 matched peaks. Furthermore, edges between two nodes were kept in the network only if each of the nodes appeared in each other’s respective top 10 most similar nodes. Finally, the maximum size of a molecular family was set to 100, and the lowest-scoring edges were removed from molecular families until the molecular family size was below this threshold. The spectra in the network were searched against GNPS spectral libraries^89^ and then were filtered in the same manner as the input data. All matches kept between network spectra, and library spectra were required to have a score above 0.65 and at least 4 matched peaks. The FBMN GNPS job is available online (https://gnps.ucsd.edu/ProteoSAFe/status.jsp?task=058516efad8f4af081b763c52404a7f7). The resulting networks were displayed and analyzed with Cytoscape (v.3.8.2)^90^.

## Acknowledgments

We are grateful to the Brazilian Biosciences National Laboratory (LNBio/CNPEM) for the use of the MS facility, and Paulo A. Baldasso for technical assistance. AD, NSM, TF and GHG thank the Fundação de Amparo à Pesquisa do Estado de São Paulo (FAPESP) grant numbers 2020/06151-4, 2018/09948-0, 2022/02992-0, 2022/03075-0 and 2021/04977-5, and the Conselho Nacional de Desenvolvimento Científico e Tecnológico (CNPq) grant numbers 311457/2020-7, 314103/2021-0 and 301058/2019-9, both from Brazil, and the National Institutes of Health/National Institute of Allergy and Infectious Diseases grant R01AI153356 (GHG) from the USA.

## Author contributions

AD and NSM conceptualized and acquired funds for the project. NSW designed and performed the majority of the experiments described, analyzed the data, and drafted the manuscript. JAG and EPA contributed to the experiments involving gene deletion and WB. GBS, AHR and NSM contributed to the expression and characterization of recombinant sirtuins. DA and TF collected and analyzed metabolomics data. CPR, FA, PAC, CFP, and GHG were responsible for in vivo experiments. LXN and AFPL designed the acetylome experiment and collected/processed the data. EV analyzed the transcriptome data. FF analyzed the cell wall component. AD and NSM supervised the project, wrote, and revised the final version of the manuscript. All authors read and approved the final version.

## Competing interest

We declare that we have no competing interests.

## SUPPLEMENTARY MATERIALS

**Sup. Table 1.**
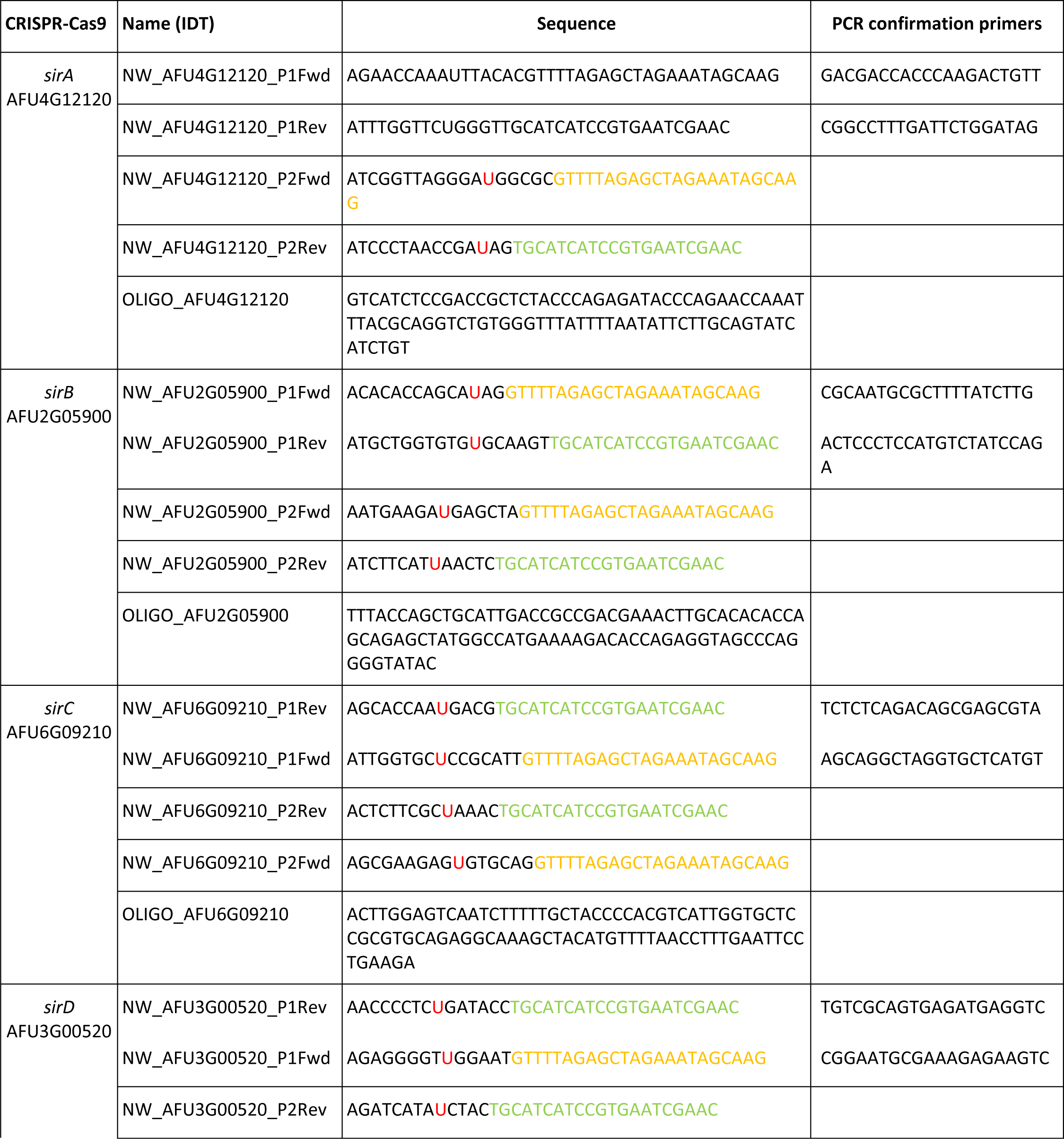
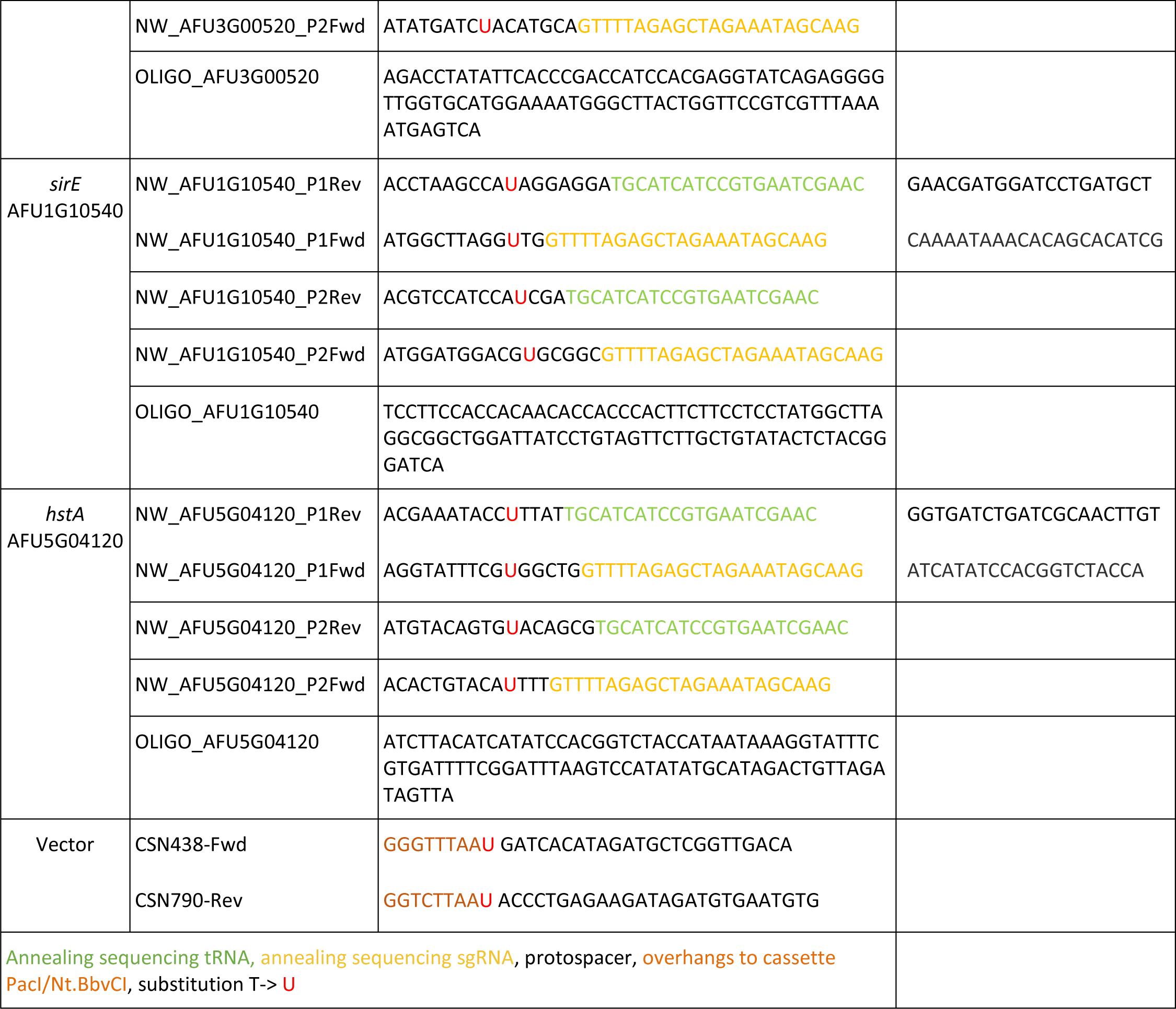
Primers and oligonucleotides used to build the biobricks for vector construction and transformation.

**Sup. Table 2.**
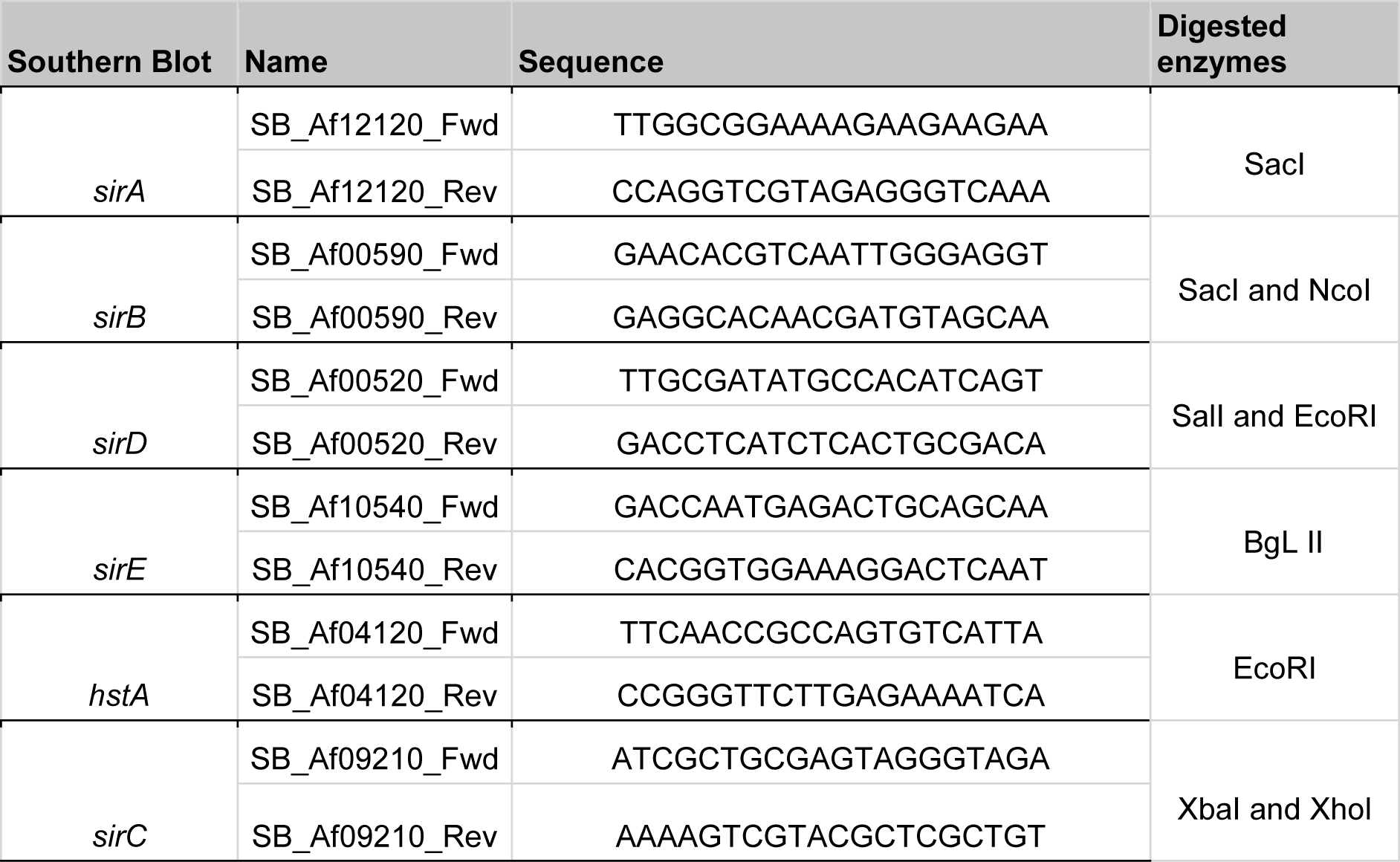
Primers used for probe sequence in southern blot assays.

**Sup. Table 3.** Total proteome of SIRTKO and ΔSirE strains.

Total proteome

**Sup. Table 4.** Acetylome analysis of Kac enriched fraction of SIRTKO and AfSirE strains.

Acetylome-Enriched fraction

**Sup. Table 5.** Genetic determinants of virulence differentially expressed in the transcriptome of SIRTKO and AfSirE strains.

RNA-seq

**Sup. Table 6.** Annotation and identification of enriched pathways using the differentially expressed genes in SIRTKO and AfSirE strains as input.

KOBAS ANALYSIS

**Sup. Figure 1.**
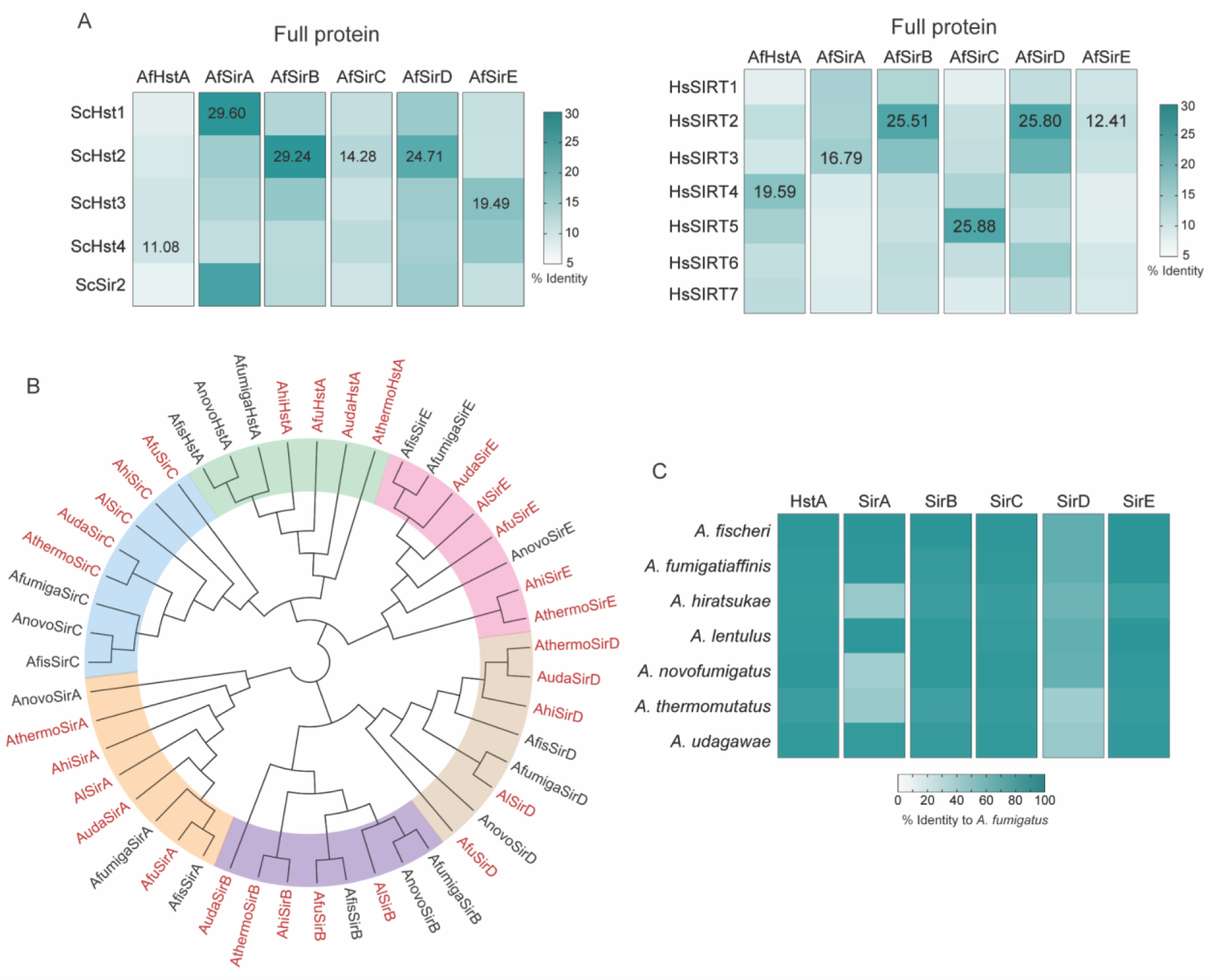
A. fumigatus sirtuins compared to other species. **A)** Amino acid identity of full-length *A. fumigatus* sirtuins compared to *S. cerevisiae* (left) and human (right) proteins. **B**. Phylogenetic tree of sirtuins from *Aspergillus* spp. Afu: *A. fumigatus*; Afi: *A. fischeri*; Afu: *A. fumigatiaffinis*; Auda: *A. udagawae*; Al: *A. lentulus*; Anovo: *A. novofumigatus*; Ahi: *A. hiratsukae*; Athermo: *A. thrmomutatus*. Species in red indicate human pathogenic species. **C**. Amino acid identity of human pathogenic *Aspergillus* species compared to *A. fumigatus*.

**Sup. Figure 2.**
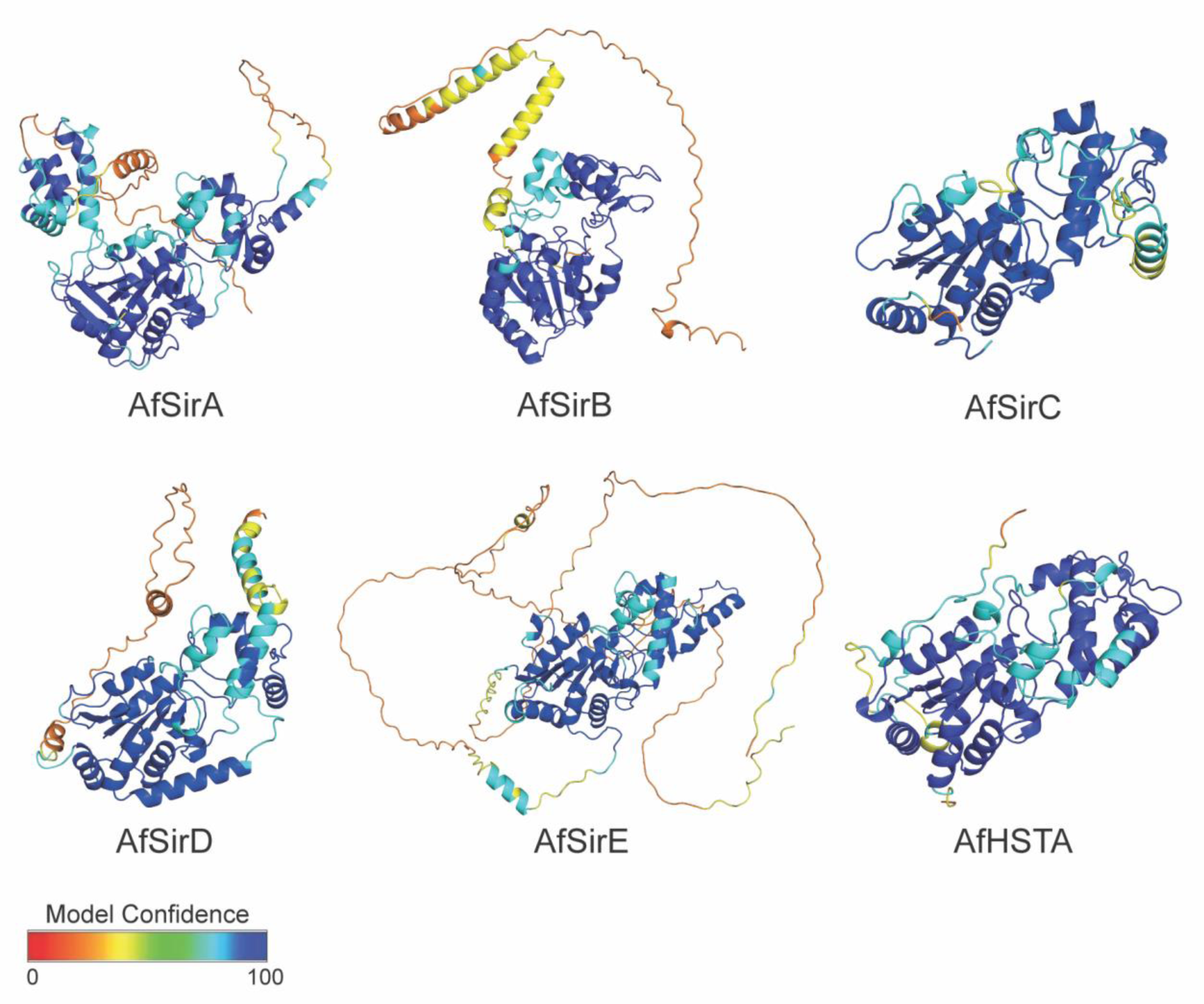
3D predicted model of *A. fumigatus* sirtuins. All models were generated using the AlphaFold tool based on the amino acid sequences deposited in the FungiDB (https://fungidb.org/fungidb/app). Dark/light blue colors indicate high confident structural regions.

**Sup. Figure 3.**
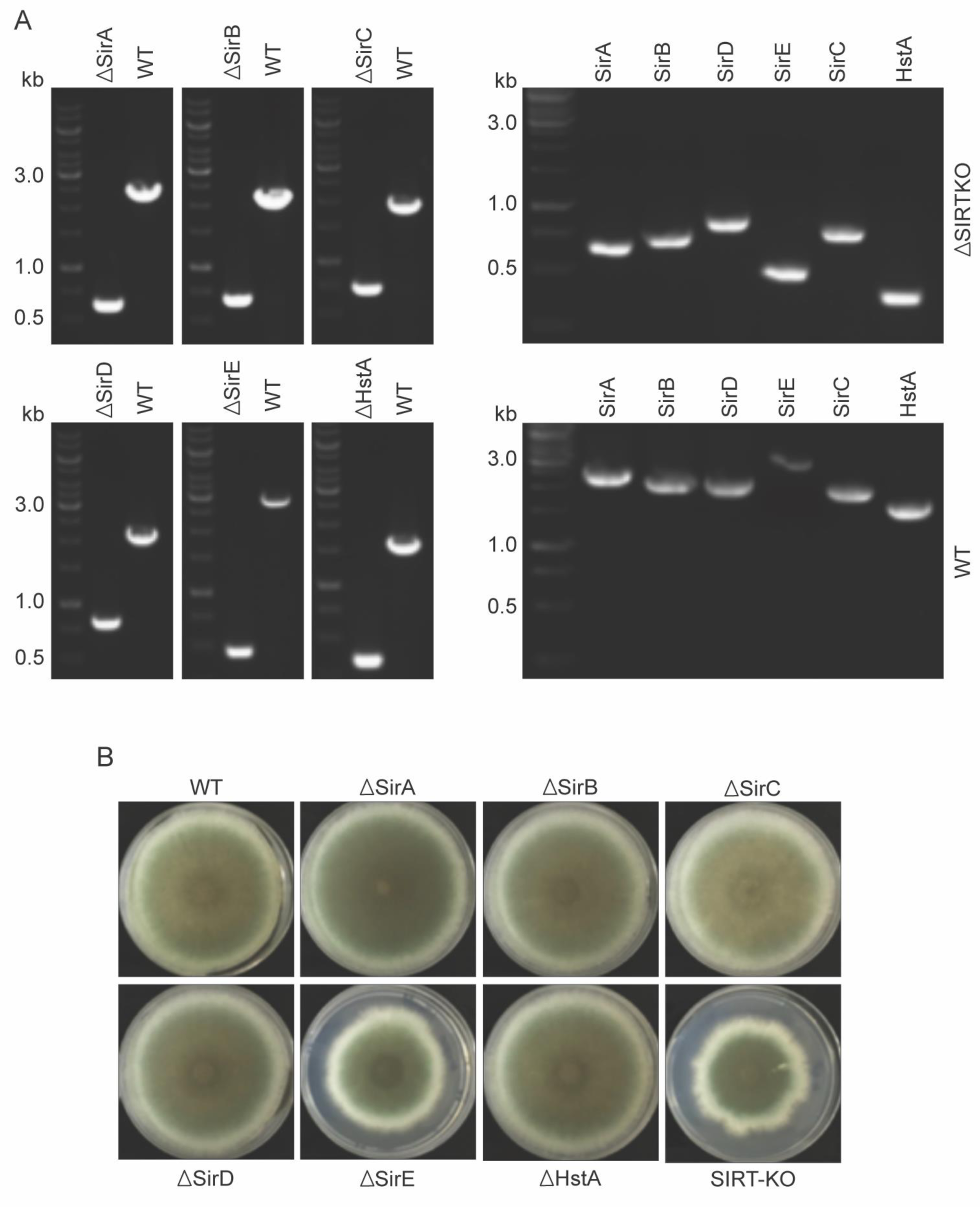
Mutants of *A. fumigatus* designed by CRISPR-Cas9 grown in Glucose Minimal Media (GMM) for 120h at 37°C and validated by diagnostic PCR confirming the deletion of the six sirtuin genes. L= ladder; Δ= mutant; WT= Wild Type.

**Sup. Figure 4.**
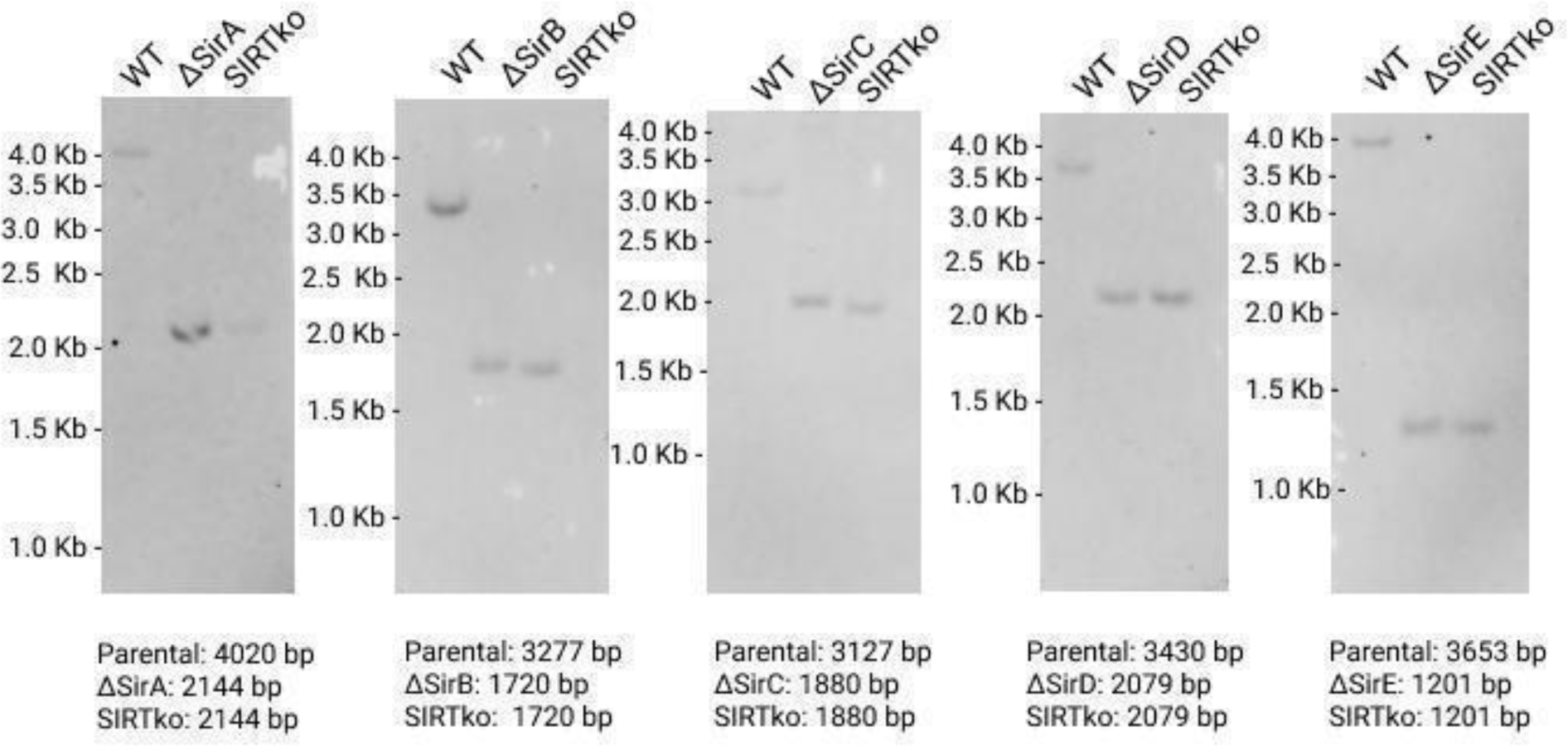
Southern blot analysis of *A. fumigatus* mutant strains. The expected fragment lengths in the single mutants and SIRTKO are detailed at the bottom of the image. Primers and digested enzymes are listed in Sup. Table 2.

**Sup. Figure 5.**
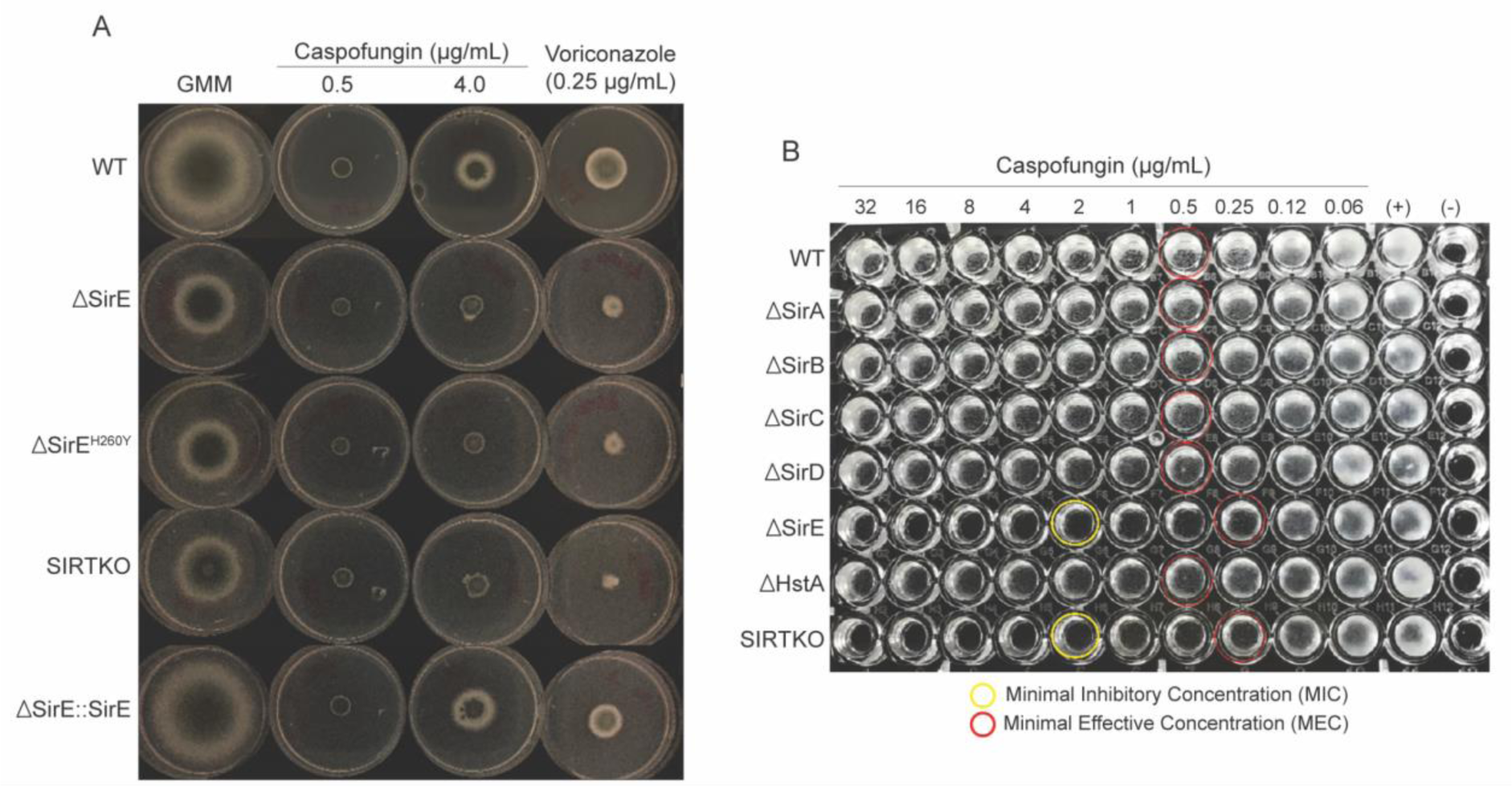
Caspofungin paradoxical effect in *A. fumigatus* strains. **A)** *A. fumigatus* sirtuin E mutant strain grown on different caspofungin concentrations and voriconazole in Glucose Minimal Media (GMM). **B)** Caspofungin Minimal Effective Concentration (MEC) analysis based on CLSI protocol (ref).

**Sup. Figure 6.**
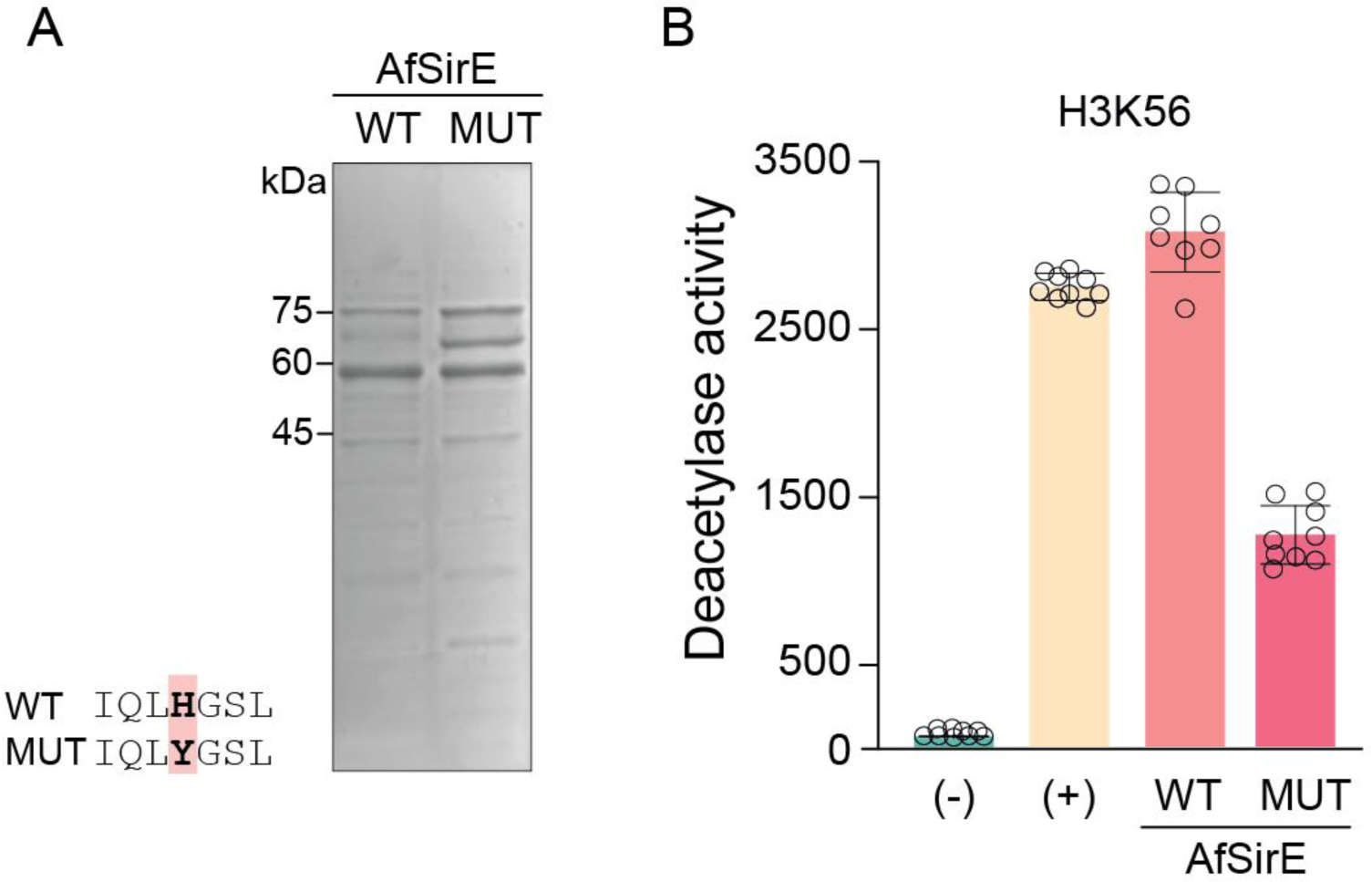
In vitro deacetylation activity of AFSirE^H260Y^ heterologous protein. **A)** SDS-PAGE of AfSirE and AfSirE^H260Y^ purified proteins used in the activity assays. **B)** Deacetylation assay using H3K56ac peptide showing the reduction in the activity of AfSirE^H260Y^ (MUT) compared to AfSirE (WT).

**Sup. Figure 7.**
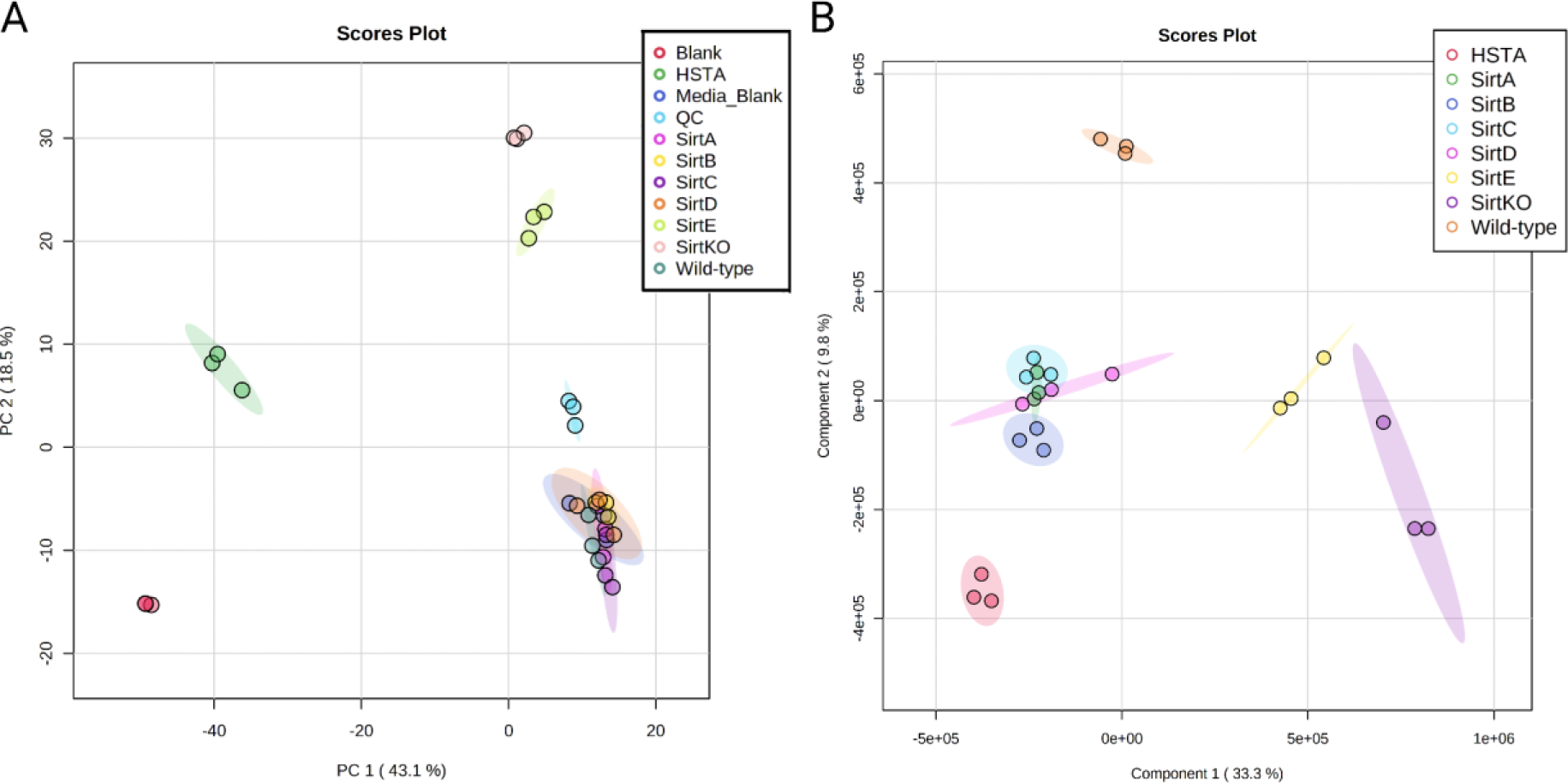
Samples distribution in the metabolome assays for secondary metabolites identification. **A**) Principal component analysis (PCA) of the methanolic extracts of *A. fumigatus* sirtuin mutant strains. PCA model built with the extracts of all biological groups, analytical blanks, QC samples and GMM media. **B)** Partial Least-squares Discriminant analysis (PLS-Da). The assay was performed in three replicates (n=3).

**Sup. Figure 8.**
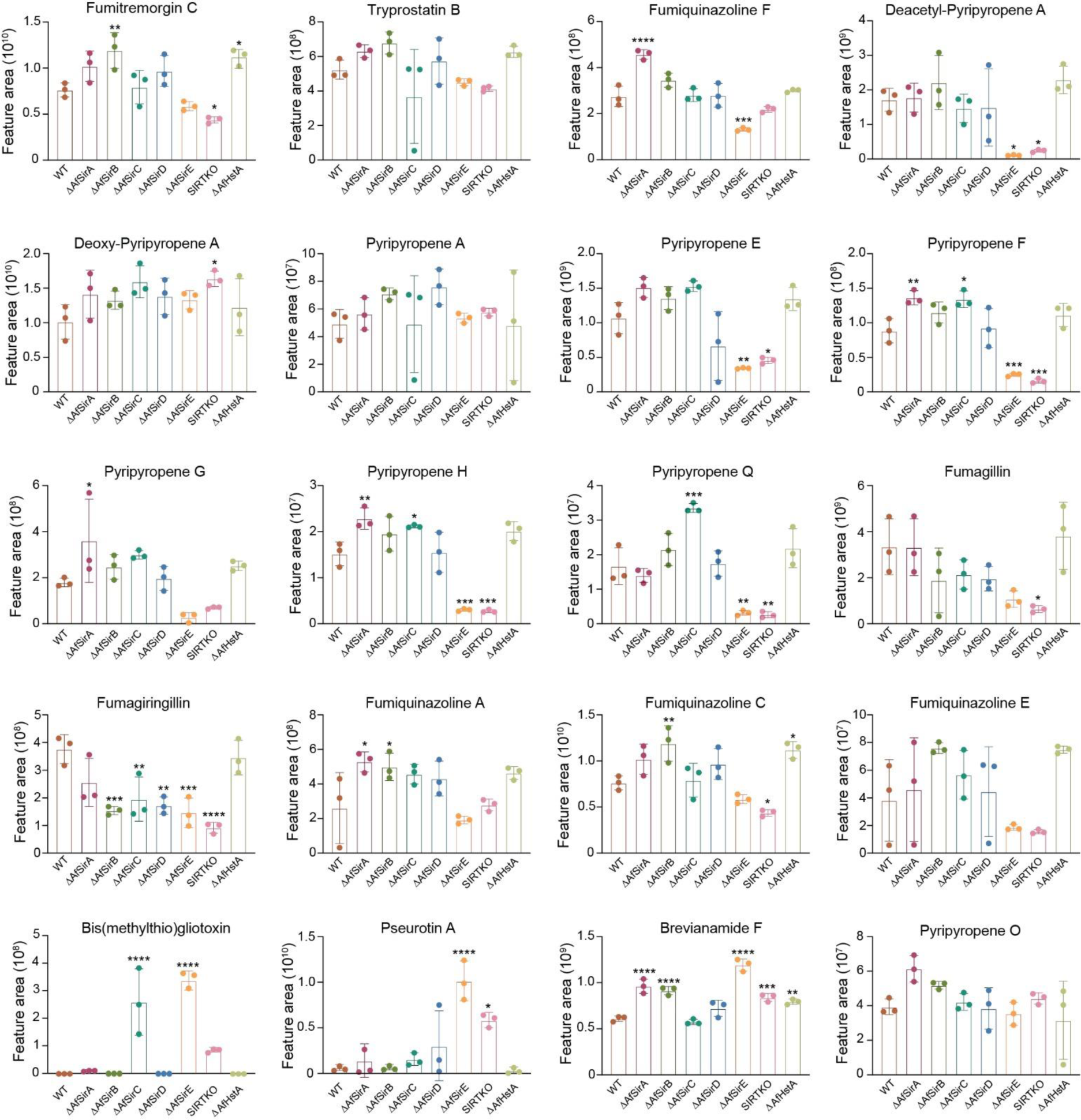
Relative quantification of putative secondary metabolites in *A. fumigatus* sirtuin mutant strains. The data represent the average value of three replicates. The error bars represent the standard deviation, p ≤ 0.05.

## Notes

### Competing Interest Statement

The authors have declared no competing interest.

## REFERENCES

1. Kwon-Chung, K. J. & Sugui, J. A. Aspergillus fumigatus—What Makes the Species a Ubiquitous Human Fungal Pathogen? PLoS Pathog 9, e1003743 (2013).

2. Latgé, J. P. & Chamilos, G. Aspergillus fumigatus and aspergillosis in 2019. Clin Microbiol Rev 33, (2020).

3. Gouzien, L. et al. Invasive Aspergillosis associated with Covid-19: A word of caution. Infect Dis Now 51, 383–386 (2021).

4. Perlin, D. S., Rautemaa-Richardson, R. & Alastruey-Izquierdo, A. The global problem of antifungal resistance: prevalence, mechanisms, and management. Lancet Infect Dis 17, e383–e392 (2017).

5. Hohl, T. M. & Feldmesser, M. Aspergillus fumigatus: Principles of pathogenesis and host defense. Eukaryotic Cell vol. 6 Preprint at 10.1128/EC.00274-07 (2007).

6. Sugui, J. A., Kwon-Chung, K. J., Juvvadi, P. R., Latge, J.-P. & Steinbach, W. J. Aspergillus fumigatus and Related Species. Cold Spring Harb Perspect Med 5, a019786– a019786 (2015).

7. Van De Veerdonk, F. L., Gresnigt, M. S., Romani, L., Netea, M. G. & Latgé, J. P. Aspergillus fumigatus morphology and dynamic host interactions. Nature Reviews Microbiology 2017 15:11 15, 661–674 (2017).

8. Chai, L. Y. A. et al. Aspergillus fumigatus Conidial Melanin Modulates Host Cytokine Response. Immunobiology 215, 915–920 (2010).

9. Rivera, A., Hohl, T. & Pamer, E. G. Immune Responses to Aspergillus fumigatus Infections. Biology of Blood and Marrow Transplantation 12, 47–49 (2006).

10. Bhabhra, R. & Askew, D. S. Thermotolerance and virulence of Aspergillus fumigatus: role of the fungal nucleolus. Med Mycol 43 Suppl 1, (2005).

11. Raffa, N. & Keller, N. P. A call to arms: Mustering secondary metabolites for success and survival of an opportunistic pathogen. PLoS Pathog 15, e1007606 (2019).

12. Sugui, J. A. et al. Gliotoxin Is a Virulence Factor of Aspergillus fumigatus: gliP Deletion Attenuates Virulence in Mice Immunosuppressed with Hydrocortisone. Eukaryot Cell 6, 1562 (2007).

13. Robbins, N., Leach, M. D. & Cowen, L. E. Lysine Deacetylases Hda1 and Rpd3 Regulate Hsp90 Function thereby Governing Fungal Drug Resistance. Cell Rep (2012) doi:10.1016/j.celrep.2012.08.035.

14. Li, W., Li, F., Zhang, X., Lin, H. & Xu, C. Insights into the post-translational modification and its emerging role in shaping the tumor microenvironment. Signal Transduction and Targeted Therapy vol. 6 Preprint at 10.1038/s41392-021-00825-8 (2021).

15. Retanal, C., Ball, B. & Geddes-Mcalister, J. Post-translational modifications drive success and failure of fungal–host interactions. Journal of Fungi vol. 7 Preprint at 10.3390/jof7020124 (2021).

16. Allfrey, V. G. & Mirsky, A. E. Structural Modifications of Histones and their Possible Role in the Regulation of RNA Synthesis. Science (1979) 144, 559–559 (1964).

17. Narita, T., Weinert, B. T. & Choudhary, C. Functions and mechanisms of non-histone protein acetylation. Nature Reviews Molecular Cell Biology (2018) doi:10.1038/s41580-018-0081-3.

18. Brownell, J. E. et al. Tetrahymena Histone Acetyltransferase A: A Homolog to Yeast Gcn5p Linking Histone Acetylation to Gene Activation. Cell 84, 843–851 (1996).

19. Downey, M. & Baetz, K. Building a KATalogue of acetyllysine targeting and function. Brief Funct Genomics 15, 109–118 (2016).

20. Taunton, J., Hassig, C. A. & Schreiber, S. L. A mammalian histone deacetylase related to the yeast transcriptional regulator Rpd3p. Science 272, 408–11 (1996).

21. Brosch, G., Loidl, P. & Graessle, S. Histone modifications and chromatin dynamics: A focus on filamentous fungi. FEMS Microbiol Rev 32, 409–439 (2008).

22. Pidroni, A., Faber, B., Brosch, G., Bauer, I. & Graessle, S. A Class 1 Histone Deacetylase as Major Regulator of Secondary Metabolite Production in Aspergillus nidulans. Front Microbiol 9, (2018).

23. Tribus, M. et al. HdaA, a Major Class 2 Histone Deacetylase of Aspergillus nidulans, Affects Growth under Conditions of Oxidative Stress. Eukaryot Cell 4, 1736–1745 (2005).

24. Tribus, M. et al. A novel motif in fungal class 1 histone deacetylases is essential for growth and development of Aspergillus. Mol Biol Cell (2010) doi:10.1091/mbc.E09-08-0750.

25. Troejer, P. et al. Histone deacetylases in fungi: Novel members, new facts. Nucleic Acids Res (2003) doi:10.1093/nar/gkg473.

26. Graessle, S. et al. Characterization of two putative histone deacetylase genes from Aspergillus nidulans. Biochimica et Biophysica Acta (BBA) - Gene Structure and Expression 1492, 120–126 (2000).

27. Izawa, M. et al. Inhibition of histone deacetylase causes reduction of appressorium formation in the rice blast fungus Magnaporthe oryzae. Journal of General and Applied Microbiology (2009) doi:10.2323/jgam.55.489.

28. Kawauchi, M. & Iwashita, K. Functional analysis of histone deacetylase and its role in stress response, drug resistance and solid-state cultivation in Aspergillus oryzae. J Biosci Bioeng 118, 172–176 (2014).

29. Kawauchi, M., Nishiura, M. & Iwashita, K. Fungus-Specific Sirtuin HstD Coordinates Secondary Metabolism and Development through Control of LaeA. Eukaryot Cell 12, 1087– 1096 (2013).

30. Ding, S. L. et al. The tig1 histone deacetylase complex regulates infectious growth in the rice blast fungus Magnaporthe oryzae. Plant Cell (2010) doi:10.1105/tpc.110.074302.

31. Lan, H. et al. The Aspergillus flavus histone acetyltransferase aflgcne regulates morphogenesis, aflatoxin biosynthesis, and pathogenicity. Front Microbiol (2016) doi:10.3389/fmicb.2016.01324.

32. Li, Y. et al. The HDF1 histone deacetylase gene is important for conidiation, sexual reproduction, and pathogenesis in Fusarium graminearum. Molecular Plant-Microbe Interactions (2011) doi:10.1094/MPMI-10-10-0233.

33. Miyamoto, A. et al. Sirtuin SirD is involved in α-amylase activity and citric acid production in Aspergillus luchuensis mut. kawachii during a solid-state fermentation process. J Biosci Bioeng 129, (2020).

34. Shwab, E. K. et al. Historie deacetylase activity regulates chemical diversity in Aspergillus. Eukaryot Cell (2007) doi:10.1128/EC.00186-07.

35. Lee, I. et al. HdaA, a class 2 histone deacetylase of Aspergillus fumigatus, affects germination and secondary metabolite production. Fungal Genetics and Biology 46, 782– 790 (2009).

36. Smith, K. M. et al. The fungus Neurospora crassa displays telomeric silencing mediated by multiple sirtuins and by methylation of histone H3 lysine 9. Epigenetics Chromatin (2008) doi:10.1186/1756-8935-1-5.

37. Itoh, E. et al. Sirtuin A regulates secondary metabolite production by Aspergillus nidulans. Journal of General and Applied Microbiology (2017) doi:10.2323/jgam.2016.11.002.

38. Itoh, E. et al. Sirtuin E is a fungal global transcriptional regulator that determines the transition from the primary growth to the stationary phase. Journal of Biological Chemistry 292, 11043–11054 (2017).

39. Wen, M. et al. Histone deacetylase SirE regulates development, DNA damage response and aflatoxin production in Aspergillus flavus. Environ Microbiol (2022) doi:10.1111/1462-2920.16198.

40. Nødvig, C. S. et al. Efficient oligo nucleotide mediated CRISPR-Cas9 gene editing in Aspergilli. Fungal Genetics and Biology (2018) doi:10.1016/j.fgb.2018.01.004.

41. Nødvig, C. S., Nielsen, J. B., Kogle, M. E. & Mortensen, U. H. A CRISPR-Cas9 system for genetic engineering of filamentous fungi. PLoS One (2015) doi:10.1371/journal.pone.0133085.

42. D’Enfert, C. et al. Attenuated virulence of uridine-uracil auxotrophs of Aspergillus fumigatus. Infect Immun 64, (1996).

43. Dauda, W. P. et al. Robust Profiling of Cytochrome P450s (P450ome) in Notable Aspergillus spp. Life 12, (2022).

44. Hokken, M. W. J. et al. Phenotypic plasticity and the evolution of azole resistance in Aspergillus fumigatus; an expression profile of clinical isolates upon exposure to itraconazole. BMC Genomics 20, (2019).

45. Huang, Z. L. et al. Interaction of a Novel Zn2Cys6 Transcription Factor DcGliZ with Promoters in the Gliotoxin Biosynthetic Gene Cluster of the Deep-Sea-Derived Fungus Dichotomomyces cejpii. Biomolecules 10, (2020).

46. Jeon, J., Kwon, S. & Lee, Y. H. Histone acetylation in fungal pathogens of plants. Plant Pathology Journal Preprint at 10.5423/PPJ.RW.01.2014.0003 (2014).

47. Li, Y. et al. Fungal acetylome comparative analysis identifies an essential role of acetylation in human fungal pathogen virulence. Commun Biol 2, 154 (2019).

48. Wassano, N. S., Leite, A. B., Reichert-lima, F. & Schreiber, A. Z. Lysine acetylation as drug target in fungi : an underexplored potential in Aspergillus spp . (2020).

49. Rack, J. G. M. et al. Identification of a Class of Protein ADP-Ribosylating Sirtuins in Microbial Pathogens. Mol Cell 59, 309–320 (2015).

50. Zhao, G. & Rusche, L. N. Genetic Analysis of Sirtuin Deacetylases in Hyphal Growth of Candida albicans. mSphere 6, (2021).

51. Ries, L. N. A. et al. Nutritional heterogeneity among aspergillus fumigatus strains has consequences for virulence in a strain- And host-dependent manner. Front Microbiol 10, (2019).

52. Latgé, J. P., Beauvais, A. & Chamilos, G. The Cell Wall of the Human Fungal Pathogen Aspergillus fumigatus: Biosynthesis, Organization, Immune Response, and Virulence. 10.1146/annurev-micro-030117-020406 71, 99–116 (2017).

53. Gastebois, A., Clavaud, C., Aimanianda, V. & Latgé, J. P. Aspergillus fumigatus: Cell wall polysaccharides, their biosynthesis and organization. Future Microbiology vol. 4 Preprint at 10.2217/fmb.09.29 (2009).

54. Valiante, V., Macheleidt, J., Föge, M. & Brakhage, A. A. The Aspergillus fumigatus cell wall integrity signalling pathway: Drug target, compensatory pathways and virulence. Front Microbiol 6, (2015).

55. Cortés, J. C. G., Curto, M. Á., Carvalho, V. S. D., Pérez, P. & Ribas, J. C. The fungal cell wall as a target for the development of new antifungal therapies. Biotechnology Advances vol. 37 Preprint at 10.1016/j.biotechadv.2019.02.008 (2019).

56. Schmalhorst, P. S. et al. Contribution of Galactofuranose to the Virulence of the Opportunistic Pathogen Aspergillus fumigatus. Eukaryot Cell 7, 1268 (2008).

57. Martín del Campo, J. S., Eckshtain-Levi, M. & Sobrado, P. Identification of eukaryotic UDP-galactopyranose mutase inhibitors using the ThermoFAD assay. Biochem Biophys Res Commun 493, 58 (2017).

58. Keller, N. P. Fungal secondary metabolism: regulation, function and drug discovery. Nature Reviews Microbiology vol. 17 Preprint at 10.1038/s41579-018-0121-1 (2019).

59. Pfannenstiel, B. T. & Keller, N. P. On top of biosynthetic gene clusters: How epigenetic machinery influences secondary metabolism in fungi. Biotechnol Adv 37, 107345 (2019).

60. Yang, K., Tian, J. & Keller, N. P. Post-translational modifications drive secondary metabolite biosynthesis in Aspergillus: a review. Environ Microbiol 24, 2857–2881 (2022).

61. Shimizu, M. et al. Hydrolase Controls Cellular NAD, Sirtuin, and Secondary Metabolites. Mol Cell Biol (2012) doi:10.1128/mcb.00032-12.

62. Shigemoto, R., Matsumoto, T., Masuo, S. & Takaya, N. 5-Methylmellein is a novel inhibitor of fungal sirtuin and modulates fungal secondary metabolite production. J Gen Appl Microbiol 64, 240–247 (2018).

63. Collemare, J. & Seidl, M. F. Chromatin-dependent regulation of secondary metabolite biosynthesis in fungi: Is the picture complete? FEMS Microbiol Rev 43, 591–607 (2019).

64. Xing, X. R. et al. Effect of nicotinamide against candida albicans. Front Microbiol 10, (2019).

65. Yan, Y., Liao, Z. Bin, Shen, J., Zhu, Z. Y. & Cao, Y. Y. Nicotinamide potentiates amphotericin B activity against Candida albicans. Virulence 13, (2022).

66. Bateman, A. et al. UniProt: the Universal Protein Knowledgebase in 2023. Nucleic Acids Res 51, (2023).

67. Jumper, J. et al. Highly accurate protein structure prediction with AlphaFold. Nature 596, (2021).

68. Varadi, M. et al. AlphaFold Protein Structure Database: Massively expanding the structural coverage of protein-sequence space with high-accuracy models. Nucleic Acids Res 50, (2022).

69. Da Silva Ferreira, M. E., et al. The akuBKU80 mutant deficient for nonhomologous end joining is a powerful tool for analyzing pathogenicity in Aspergillus fumigatus. Eukaryot Cell 5, (2006).

70. Nørholm, M. H. H. A mutant Pfu DNA polymerase designed for advanced uracil-excision DNA engineering. BMC Biotechnol (2010) doi:10.1186/1472-6750-10-21.

71. Nour-Eldin, H. H., Hansen, B. G., Nørholm, M. H. H., Jensen, J. K. & Halkier, B. A. Advancing uracil-excision based cloning towards an ideal technique for cloning PCR fragments. Nucleic Acids Res (2006) doi:10.1093/nar/gkl635.

72. Geu-Flores, F., Nour-Eldin, H. H., Nielsen, M. T. & Halkier, B. A. USER fusion: A rapid and efficient method for simultaneous fusion and cloning of multiple PCR products. Nucleic Acids Res (2007) doi:10.1093/nar/gkm106.

73. Jin, F. J., Katayama, T., Maruyama, J. ichi & Kitamoto, K. Comparative genomic analysis identified a mutation related to enhanced heterologous protein production in the filamentous fungus Aspergillus oryzae. Appl Microbiol Biotechnol (2016) doi:10.1007/s00253-016-7714-2.

74. Maniatis, T., Fritsch, E. F. & Sambrook, J. Molecular Cloning : a laboratory manual. 2nd edition. Cold Spring Harbor, New York 0, (1988).

75. François, J. M. A simple method for quantitative determination of polysaccharides in fungal cell walls. Nature Protocols 2007 1:6 1, 2995–3000 (2007).

76. CLSI. Reference Method for Broth Dilution Antifungal Susceptibility Testing of Filamentous Fungi ; Approved Standard — Second Edition. vol. 28 (2008).

77. Fuchs, B. B., O’Brien, E., Khoury, J. B. E. & Mylonakis, E. Methods for using Galleria mellonella as a model host to study fungal pathogenesis. Virulence 1, (2010).

78. Brauer, V. S., et al. Extracellular Vesicles from Aspergillus flavus Induce M1 Polarization In Vitro. mSphere 5, (2020).

79. Ewels, P., Magnusson, M., Lundin, S. & Käller, M. MultiQC: summarize analysis results for multiple tools and samples in a single report. Bioinformatics 32, 3047–3048 (2016).

80. Bolger, A. M., Lohse, M. & Usadel, B. Trimmomatic: a flexible trimmer for Illumina sequence data. Bioinformatics 30, 2114–2120 (2014).

81. Dobin, A. et al. STAR: Ultrafast universal RNA-seq aligner. Bioinformatics 29, (2013).

82. Liao, Y., Smyth, G. K. & Shi, W. FeatureCounts: An efficient general purpose program for assigning sequence reads to genomic features. Bioinformatics 30, (2014).

83. Robinson, M. D., McCarthy, D. J. & Smyth, G. K. edgeR: A Bioconductor package for differential expression analysis of digital gene expression data. Bioinformatics 26, (2009).

84. Bradford, M. M. A Rapid and Sensitive Method for the Quantitation Microgram Quantities of Protein Utilizing the Principle of Protein-Dye Binding. 254, 248–254 (1976).

85. Schilling, B., Meyer, J. G., Wei, L., Ott, M. & Verdin, E. High-Resolution Mass Spectrometry to Identify and Quantify Acetylation Protein Targets. Methods Mol Biol 1983, 3 (2019).

86. Chambers, M. C., et al. A cross-platform toolkit for mass spectrometry and proteomics. Nature Biotechnology vol. 30 Preprint at 10.1038/nbt.2377 (2012).

87. Nothias, L. F. et al. Feature-based molecular networking in the GNPS analysis environment. Nat Methods 17, (2020).

88. Chong, J., Wishart, D. S. & Xia, J. Using MetaboAnalyst 4.0 for Comprehensive and Integrative Metabolomics Data Analysis. Curr Protoc Bioinformatics 68, (2019).

89. Wang, M., et al. Sharing and community curation of mass spectrometry data with Global Natural Products Social Molecular Networking. Nature Biotechnology vol. 34 Preprint at 10.1038/nbt.3597 (2016).

90. Shannon, P. et al. Cytoscape: A software Environment for integrated models of biomolecular interaction networks. Genome Res 13, (2003).

